# Establishment of pan-Influenza A (H1-H18) and pan-Influenza B (pre-split, Vic/Yam) Pseudotype Libraries for efficient vaccine antigen selection

**DOI:** 10.1101/2021.05.20.444964

**Authors:** Joanne Marie M. Del Rosario, Kelly A.S. da Costa, Benedikt Asbach, Francesca Ferrara, Matteo Ferrari, David A. Wells, Gurdip Singh Mann, Veronica O. Ameh, Claude T. Sabeta, Ashley C. Banyard, Rebecca Kinsley, Simon D. Scott, Ralf Wagner, Jonathan L. Heeney, George W. Carnell, Nigel J. Temperton

**Affiliations:** Viral Pseudotype Unit, Medway School of Pharmacy, The Universities of Greenwich and Kent at Medway, Chat-ham, United Kingdom; Department of Physical Sciences and Mathematics, College of Arts and Sciences, University of the Philippines Manila, Manila, Philippines; DIOSynVax, Cambridge, United Kingdom; Institute of Medical Microbiology and Hygiene, University of Regensburg, Regensburg, Germany; Department of Veterinary Medicine, University of Cambridge, Cambridge, United Kingdom; Federal University of Agriculture Makurdi, College of Veterinary Medicine, Department of Veterinary Public Health and Preventive Medicine, P.M.B. 2373, Makurdi Bene State, Nigeria; University of Pretoria, Faculty of Veterinary Science, Department of Veterinary Tropical Diseases, P. Bag X04, Onderstepoort, South Africa; OIE Rabies Reference Laboratory, Agricultural Research Council- Onderstepoort Veterinary Research, Onderstepoort, South Africa; Animal and Plant Health Agency (APHA), Department of Virology, Weybridge, Surrey, United Kingdom; Institute of Clinical Microbiology and Hygiene, University Hospital Regensburg, Regensburg, Germany

**Keywords:** influenza, hemagglutinin, pseudotype, vaccine, immunogenicity, monoclonal antibody, neutralization

## Abstract

We have developed an influenza hemagglutinin (HA) pseudotype library encompassing Influenza A subtypes HA1-18, and Influenza B subtypes (both lineages) to be employed in influenza pseudotype microneutralization (pMN) assays. The pMN is highly sensitive and specific for detecting virus-specific neutralizing antibodies against influenza viruses and can be used to assess antibody functionality *in vitro*. Here we show the production of these viral HA pseudotypes and their employment as substitutes for wildtype viruses in influenza serological and neutralization assays. We demonstrate its utility in detecting serum response to vaccination with the ability to evaluate cross-subtype neutralizing responses elicited by specific vaccinating antigens. Our findings may inform further pre-clinical studies involving immunization dosing regimens in mice and may help in the creation and selection of better antigens for vaccine design. These HA pseudotypes can be harnessed to meet strategic objectives that contribute to the strengthening of global influenza surveillance, expansion of seasonal influenza prevention and control policies, and strengthening pandemic preparedness and response.

## 1. Introduction

Influenza viruses are segmented, negative sense, single-stranded, enveloped RNA viruses belonging to the Orthomyxoviridae family [1, 2]. Within this family, there are three types of influenza virus that circulate in humans, influenza A, B, and C [3-5]. Only influenza A (IAV) and influenza B (IBV) viruses are endemic in the global human population, rapidly spreading around the world in seasonal epidemics; imposing considerable economic burden and death [6, 7]. From its wild bird reservoir, IAV is able to transmit from domestic poultry [8], which is the gateway to infection of mammals, most notably, swine and humans [9]. IBV’s natural reservoir is humans, however there have been reports of infection in seals [10, 11], alluding to its potential to cause disease in other species.

Influenza A, and to a lesser extent, influenza B can be further classified by structural and genetic differences in the two most abundant glycoproteins expressed on the viral surface – hemagglutinin (HA) which is required for viral entry and fusion [12-14], and neuraminidase (NA), which is involved in release of viral progeny [15]. Currently, 18 distinct antigenic HA (H1-H18) and 11 antigenic NA (N1-N11) subtypes have been described for IAV [5, 6, 16]. Based on phylogenetic analysis, IAV HA subtypes are divided into two groups: Group 1 - H1, H2, H5, H6, H8, H9, H11, H12, H13, H16, H17 and H18 subtypes, and Group 2 - H3, H4, H7, H10, H14 and H15 [15]. IBV is not as diverse and has consequently been divided into 2 distinct lineages, B/Yamagata-like and B/Victoria-like viruses [17].

Hemagglutinin is a trimeric glycoprotein consisting of a globular head attached to a fibrous stem [14, 18, 19]. The HA head is highly antigenic and is subject to mutations and re-assortment of genetic material over time [9, 20, 21]. Minor genetic changes such as single point mutations in the HA head are known to give rise to antigenic drift. In contrast, antigenic shift, wherein an influenza A virus strain acquires an HA or NA segment from another subtype of IAV usually from a zoonotic reservoir can also occur leading to the emergence of new variants or strains [20, 22-27]. Antigenic shift is of concern as it may result in the emergence of completely novel virus to which the human population has no pre-existing immunity, and as such, may have pandemic potential. To date only 3 HA (H1, H2 & H3) and 2 NA (N1 & N2) subtypes are known to have caused human pandemics [28-31].However this does not preclude other subtypes becoming pandemic in the human population in the future and as such, the emergence of novel IAV strains remains a major concern. There have been numerous documented cases of human infection with highly pathogenic influenza A viruses (HPAI) H5 and H7, viral subtypes that predominantly cause outbreaks in poultry [32-35]. Nonetheless, these incidences have not yet resulted in these viruses acquiring the ability to sustain human to human transmission [36-38]. Whilst antigenic divergence both within and across HA subtypes exists,the HA stem domain is more conserved and although not as immunogenic as the head domain [15, 20], is increasingly being explored as a candidate for universal influenza vaccines [20, 39]. As such, the importance of studying HA structure and function and monitoring antigenic changes within HA is critical to: understanding antigenic evolution; defining the most antigenically relevant antigens for annual human vaccination programs [40, 41], determining potent universal vaccine targets [42, 43], developing vaccines for veterinary use [8, 44], and improving influenza diagnosis and therapeutic interventions [45-48].

Vaccine strain selection for seasonal influenza is carried out via the hemagglutinin inhibition (HI) assay that antigenically characterizes influenza viruses [49, 50]. The HI test works by measuring the interaction between serum antibody and the influenza HA domain of currently circulating IAV and IBV strains and the resulting inhibition of red blood cell agglutination and is currently the measure for seroconversion and protection [41, 51-53]. To improve current vaccination strategies and aid the development of a universal influenza vaccine, additional reliable tools are necessary to identify and progress promising candidates specifically targeting both the hemagglutinin head and stem domains [47, 54-56]. The advent of pseudotyped lentiviral vectors have enabled the study of HA interactions with antibodies, drugs and host cell receptors with ease [57-59]. These pseudotypes undergo abortive replication and do not give rise to replication-competent progeny [60, 61]. While it is logistically possible to deal with low pathogenic strains of influenza, studies on strains that are exotic and not widespread in the population are considerably hampered by availability of BSL facilities and highly trained and qualified personnel required for handling and processing these viruses.

To address these issues, we have constructed a comprehensive library of IAV and IBV HA pseudotypes that we tested against available antisera, HA stem-directed monoclonal antibodies, and to detect neutralizing responses in sera in mouse vaccine studies to produce optimized seasonal vaccines and candidate pandemic vaccines. This repository of pseudotypes is contributing to the World Health Organization’s global influenza strategy for 2019-2030 of “ Prevent, Control and Prepare” [62], with the goal of employing these PV as tools to further vaccine R&D that will contribute to reducing the burden of seasonal influenza, minimizing the risk of zoonotic influenza, and mitigating the impact of pandemic influenza.

## 2. Materials and Methods

### 2.1. Plasmid production and transformation

Hemagglutinin genes from Influenza A virus (IAV) subtypes HA1-18 and Influenza B (IBV) B/Victoria-like and B/Yamagata-like viruses were cloned in either pI.18 (in house), phCMV1 (GenScript, Netherlands), or pEVAC plasmids (GeneArt, Germany). pI.18 is a high-copy Amp^R^ pUC-based plasmid that permits robust mammalian gene expression in various cell types via the human cytomegalovirus (hCMV) immediate-early gene promoter and the enhancer hCMV Intron A [63]. phCMV1 (Genlantis) is a constitutive mammalian gene expression vector driven by a modified hCMV immediate-early promoter and enhancer/intron together with a Simian Vacuolating virus 40 (SV40) promoter. Kan^R^ and Neo^R^ allow selection of plasmid-positive prokaryotic and eukaryotic cells, respectively in phCMV. pEVAC is also a mammalian expression vector with an hCMV immediate-early promoter/enhancer followed by an intron (HTLV-1-R splice donor and hCMV-IE splice acceptor), a BGH poly-adenylation sequence and Kan^R^ gene. All HA genes were gene-optimized and adapted to human codon use using the GeneOptimizer algorithm [64] and have a strong Kozak-initation motif.

Influenza hemagglutinin plasmid constructs were generated by cloning the IAV or IBV HA transgenes into pI.18, phCMV1, or pEVAC via restriction digest into the plasmids’ multiple cloning site (MCS). Plasmids were transformed in chemically induced competent *E. coli* DH5α cells (Invitrogen 18265-017) via the heat-shock method. Plasmid DNA was recovered from transformed bacterial cultures via the Plasmid Mini Kit (Qiagen 12125) or the endotoxin-free HiSpeed Plasmid Midi Kit (Qiagen 12643). All DNA extracts were quantified using UV spectrophotometry (NanoDrop™ -Thermo Scientific).

### 2.2. Propagation and maintenance of cell cultures

Human Embryonic Kidney (HEK) 293T/17 (ATCC: CRL-11268^a^) cells were used for production and titration of pseudotyped lentiviral vectors and neutralization assays. Madin-Darby Canine Kidney (MDCK) II cells were used for titration and neutralization assays of Influenza H17 and H18 pseudotyped viruses. Both cell lines were maintained in complete medium, Dulbecco’s Modified Essential Medium (DMEM) (PANBiotech P04-04510) with high glucose and GlutaMAX. DMEM was supplemented with 10% (v/v) heat-inactivated Fetal Bovine Serum (PANBiotech P30-8500), and 1% (v/v) Penicillin-Streptomycin (PenStrep) (Sigma P4333). Cells were incubated at 37°C and 5% CO_2_.

### 2.3. Transfection of influenza HA pseudotypes (PV)

Influenza HA pseudotypes were produced as described previously [58]. Briefly, 4×10^5^ HEK 293T/17 cells in complete DMEM were seeded per well of a 6-well plate and incubated at 37°C, 5% CO_2_ overnight. The next day, media was replaced and cells were transfected using Opti-MEM™ (Thermo Fisher Scientific 31985062) and FuGENE® HD Transfection Reagent (ProMega E2312) with the following plasmids: HA encoding plasmid (pI.18/phCMV1/pEVAC), luciferase reporter plasmid pCSFLW [57], and p8.91 gag-pol (Gag-Pol expression plasmid [60, 61]). Plates were incubated at 37°C, 5% CO_2_. For transfection of low pathogenicity avian influenza (LPAI) and other subtypes with a monobasic cleavage site, an additional plasmid expressing either type II Transmembrane Protease Serine 2 (TMPRSS2) [65], type II Transmembrane Protease Serine 4 (TMPRSS4) [66], or human airway trypsin-like protease (HAT) [65], was also included. For the H18 subtype, 50 ng of A/flat-faced bat/Peru/033/2010/N11 in pEVAC was also included. Amounts of plasmid DNA and reagents used for transfection in a single well of a 6-well plate are indicated in **Table 1**. All plasmid DNA were combined in OptiMEM and FuGENE® HD added dropwise followed by incubation for 15 minutes. The plasmid DNA-OptiMEM mixture was then added to the cells with constant swirling. At least 8 hours post-transfection, 1 unit of exogenous Neuraminidase (Sigma N2876) was added to the 6 well-plates, with the exception of the H18 subtype. Forty-eight hours post-transfection, supernatants were collected, passed through a 0.45 μm filter and stored at -80°C.

**Table 1.**
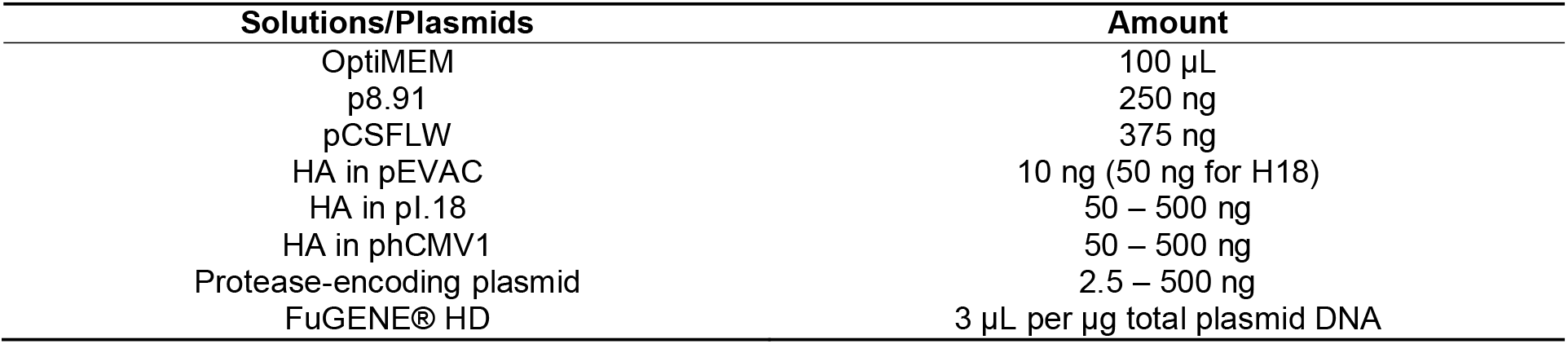
Amounts of Influenza HA transfection components.

### 2.4. Influenza pseudotype titration

Titration experiments were performed in Nunc F96 MicroWell white opaque polystyrene plates (Thermo Fisher Scientific 136101). The pseudotype production titer was evaluated by transducing HEK293T/17 cells (or MDCKII cells for H17 and H18) with the PV. Fifty microliters of viral supernatant were serially diluted two-fold across a 96-well plate in duplicate before adding 50 μL of 1⨯10^4^ HEK293T/17 cells to each well. Control wells in which there was no PV added were also present on each plate as an indirect cell viability measurement. Plates were then incubated at 37°C, 5% CO_2_ for 48 hours. Media was removed and 25 µL Bright-Glo® luciferase assay substrate was added to each well. Titration plates were then read using the GloMax® Navigator (ProMega) using the Promega GloMax® Luminescence Quick-Read protocol. Viral pseudotype titer was then determined in Relative Luminescence Units/mL (RLU/mL).

### 2.5. Reference antisera and bat surveillence sera

Reference antisera to assess the neutralization sensitivity of representative IAV and IBV pseudotypes from our library was obtained from the OIE (World Organisation for Animal Health), the National Institute for Biological Standards and Control (NIBSC) or the Animal and Plant Health Agency (APHA). Antisera were generated by immunizing chickens (OIE) and sheep (NIBSC) with HA antigen. At the time of publication, reference antiserum for H17 was not available, however, Fugivorous bat sera, collected as part of a bat sera surveillance program in Nigeria, was provided by APHA.

### 2.6. Mouse immunogenicity studies

For mouse immunogenicity studies, 6-8 week old female BALB/c mice were obtained from Charles River Laboratories and housed at University Biomedical Services, University of Cambridge. Mice were divided into groups of six for each individual vaccination antigen. On day 0, mice were injected subcutaneously (SC) on the rear flank with a 50 μL volume of 50 µg pEVAC HA, produced using the EndoFree Plasmid Mega Kit (Qiagen), or PBS for negative control groups. Immunizations were repeated on weeks 2, 4, and, 6 (**Figure 1**). Mice were weighed daily and monitored for any signs of disease or distress. Mice were bled at 42 days post immunization (dpi), 56 dpi, and 70 dpi (**Figure 1**). 70 days post immunization, all mice were culled and terminal bleeds collected. Collected blood was left to clot for 1 hour at room temperature and serum was separated via centrifugation at 2,000xg for 10 minutes at 4°C and stored at -20°C.

**Figure 1.**
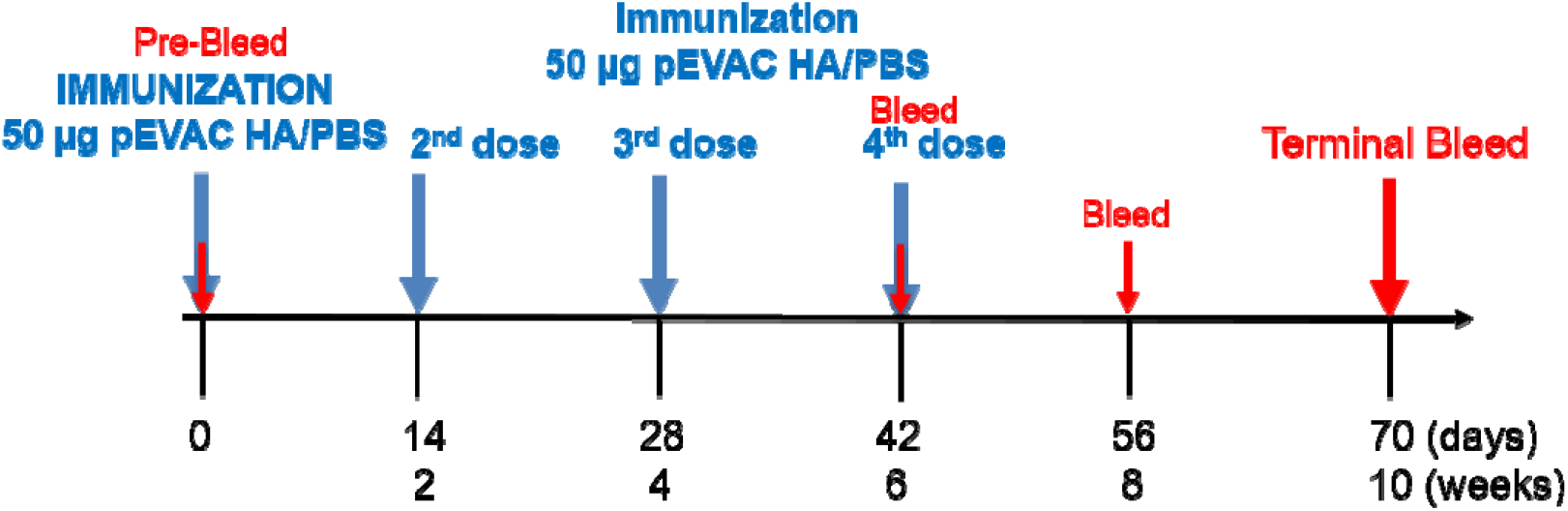
Study schedule of immunization with pEVAC HA antigens. Mice received either pEVAC HA antigens or PBS (negative control groups) on weeks 0, 2, 4, and 6 via subcutaneous rear flank injection. Blood was collected on weeks 6, 8, and 10.

### 2.7. Pseudotype microneutralization (pMN) assay

We performed pseudotype microneutralization assays using standard reference antisera, monoclonal antibodies (mAb), and serum samples from animal studies. Monoclonal antibody concentrations used were in the range of 0.5 ng/mL - 1000 ng/mL and serum and antiserum samples were initially diluted 1:20 or 1:50 in 50 µL complete DMEM, before being serially diluted two-fold across a 96-well plate. Fifty microliters of PV at a concentration of 1.0⨯10^6^ RLU/well as determined via titration was then added to the mAb or serum dilutions, making the final dilution of sera 1:40 or 1:100. This mixture was incubated for 1 hour at 37°C, 5% CO_2_. Afterwards, 50 µL of 1.5⨯10^4^ HEK293T/17 cells were added to each well. PV only (equivalent to 0% neutralization) and cell only controls with no virus (equivalent to 100% neutralization control) were also included in the test plate. Plates were incubated for 48 hours at 37°C and 5% CO_2_. Media was removed and 25µL Bright-Glo® luciferase assay substrate added to each well. Plates were then read using the GloMax® Navigator (ProMega) using the Promega GloMax® Luminiscence Quick-Read protocol. Half-maximal inhibitory dilution or concentration (IC_50_) values were calculated using GraphPad Prism 8.12. A detailed analysis is described in Ferrara, 2018 [67].

### 2.8. Statistical analysis

All statistical analyses were performed with GraphPad Prism 8.12 for Windows (GraphPad Software). The Kruskal-Wallis H test, a rank-based nonparametric test, was used to determine if there were statistically significant differences between two or more groups in comparison to a control group.

### 2.9. Bioinformatic analysis

HA sequences for both IAV and IBV were downloaded from the Influenza Virus Resource database (IVRD) (fludb.org). Phylogenetic tree was generated using the Cyber-Infrastructure for Phylogenetic RE-Search (CIPRES) Gateway [68]. The resulting tree file was then visualized using the Archaeopteryx tree viewer in the Influenza Resource Database (IRD) [69].

## 3. Results

### 3.1. Production of IAV and IBV Pseudotype library

The influenza pseudotype viruses (PV) described herein were constructed using the transfection method detailed above (Section 2.3). All PV were produced with the following 3 plasmids: i) a plasmid containing packaging genes from a surrogate lentivirus (HIV) (gag-pol) which is defective for native HIV envelope, ii) a plasmid expressing the HA envelope of the strain being studied (IAV or IBV), and iii) a transfer plasmid expressing firefly luciferase reporter (**Figure 2a**). 1 unit of exogenous neuraminidase (exoNA) was added per well to facilitate viral egress, with the PV containing the HA envelope on its surface, harvested in cell supernatants.

**Figure 2.**
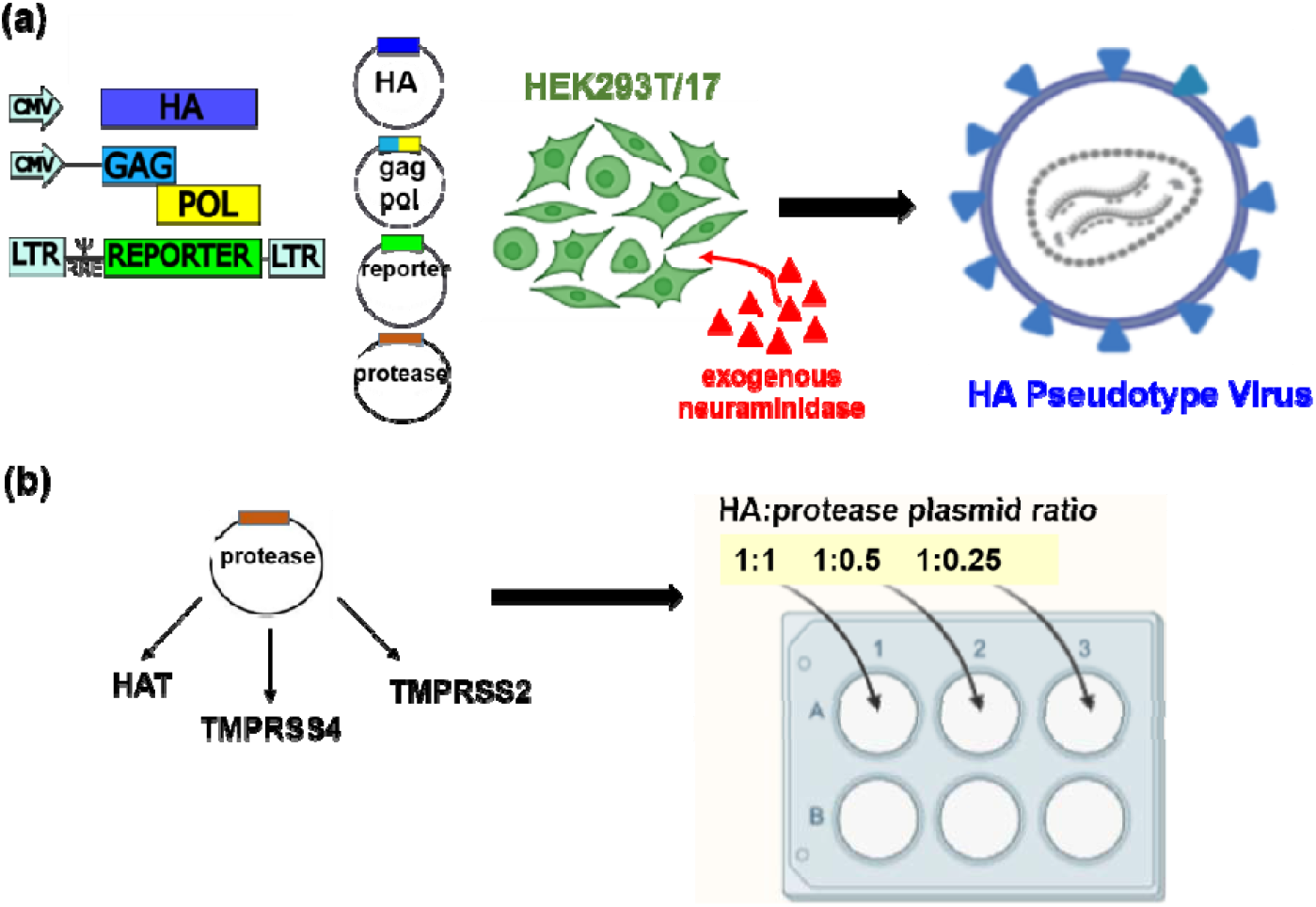
Schematic representation of the production of influenza HA pseudotypes by plasmid transfection. **(a)** Using the 3 or 4 plasmid system, exogenous NA was added to the transfected cultures at least 8 hours post-transfection. **(b)** Addition of protease, which is necessary for production of low pathogenic influenza A viruses (LPAI) and subtypes with a monobasic cleavage site, and IBV, were optimized to increase titers by transfecting in a ‘checkerboard’ approach with different proteases (e.g. HAT, TMPRSS4 & TMPRSS2). Protease plasmid was added at a ratio of 1:1, 1:0.5 and 1:0.25 to HA plasmid DNA for rapid optimization in a 6 well plate format. All pseudotypes were harvested after 48 hours in culture. Image created in BioRender.

IAV and IBV strains which contain monobasic cleavage sites require the presence of a trypsin-like protease *in vitro* to catalyze HA proteolytic cleavage from the inactive trimeric HA0 to the active HA1 and HA2 leading to viral membrane fusion [70-72]. As demonstrated previously, an additional plasmid expressing a trypsin-like protease was required for PV production (**Figure 2a**) [58, 65, 66], with the amount of protease plasmid DNA requiring optimization for each PV produced. We found that this is dependent on the HA subtype and occasionally the strain being produced (**Figure 3, Table 3**). Optimization was achieved using a 6-well plate checkerboard system for protease amounts (**Figure 2b**), and a fixed amount of all other plasmids was used to transfect 293T/17 cells. For all strains except HPAI strains, initial transfections were under-taken with human airway trypsin-like protease (HAT) in the top 3 wells and Transmembrane Serine Protease 4 (TMPRSS 4) in the bottom 3 wells of the 6-well plate. Protease plasmid was added at a ratio to the HA plasmid (**Figure 2b**), i.e., for every 10 ng of HA, we tested with 10 ng, 5 ng, and 2.5 ng protease DNA. All PV produced were titrated and PV titer determined in RLU (**Figure 3**).

**Figure 3.**
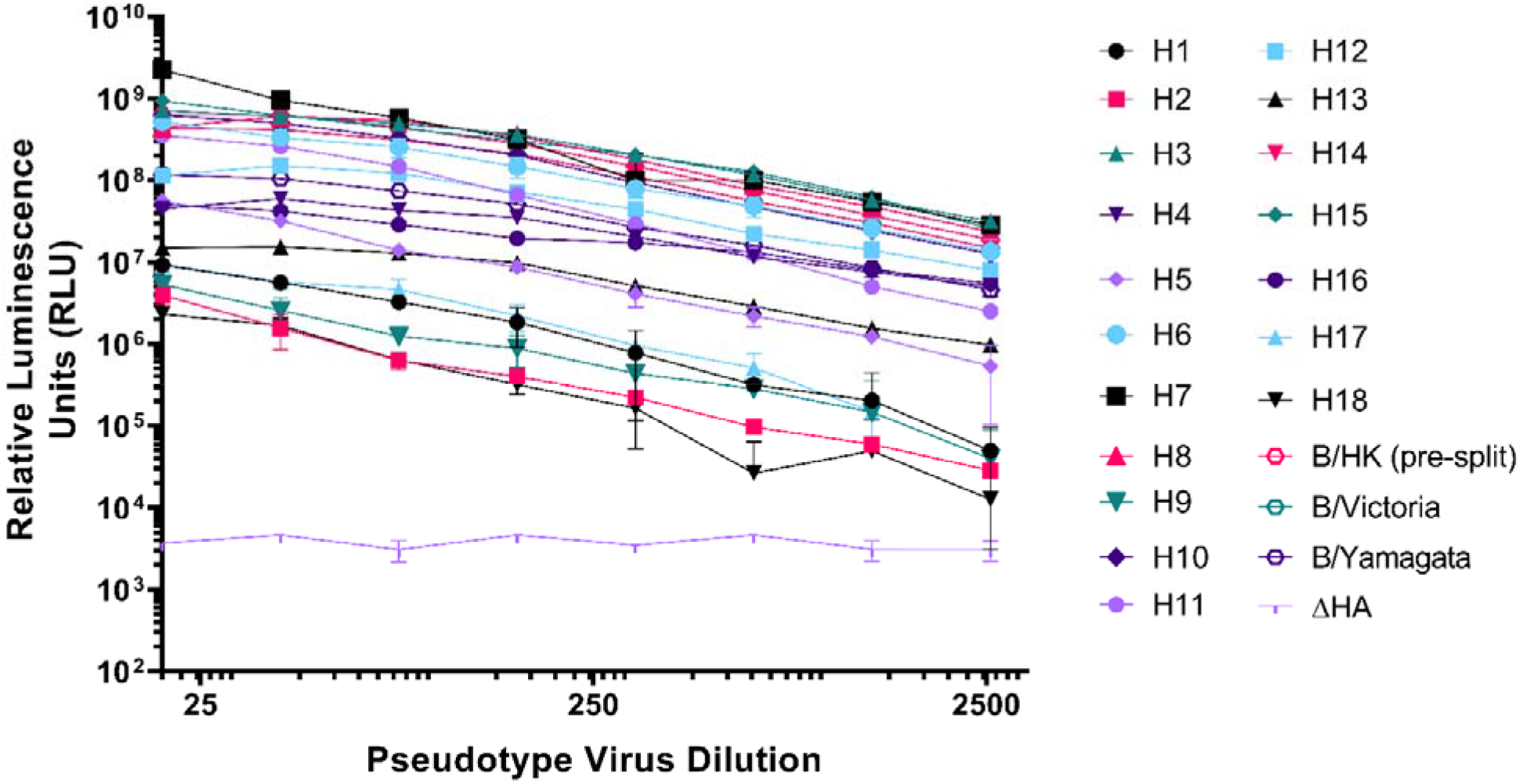
Titration of representative Influenza A (H1-H18) and influenza B (pre-split, B/Victoria-like and B/Yamagata-like lineages) viruses. Pseudotyped lentiviral particles with HA envelopes: H1 – A/England/195/2009(H1), H2 – A/quail/Rhode Island/16-0186222-1/2016(H2), H3 – A/ruddy turnstone/Delaware Bay/606/2017(H3), H4 – A/green-winged teal/California/K218/2005(H4), H5 – A/gyrfalcon/Washington/41088-6/2014(H5), H6 – A/American wigeon/California/HS007A/2015(H6), H7 – A/Shanghai/2/2013(H7), H8 – A/mallard/Netherlands/7/2015(H8), H9 – A/chicken/Israel/291417/2017(H9), H10 – A/duck/Bangladesh/24268/2015(H10), H11 – A/red shoveler/Chile/C14653/2016(H11), H12 – A/northern shoveler/Nevada/D1516557/2015(H12), H13 – A/laughing gull/New Jersey/UGAI7-2843/2017(H13), H14 – A/mallard/Astrakhan/263/1982(H14), H15 – A/duck/Bangladesh/24697/2015(H15), H16 – A/black-headed gull/Netherlands/1/2016(H16), pre-split – B/Hong Kong/8/1973, B/Victoria – B/Brisbane/60/2008, and B/Yamagata – B/Phuket/3073/2013, were titrated in HEK293T/17 cells. H17 – A/little yellow-shouldered bat/Guatemala/60/2017(H17), and H18 – A/flat-faced bat/Peru/33/2010(H18) were titrated in MDCKII cells. ΔHA is included as a no envelope control. Each point represents the mean and standard deviation of two replicates per dilution. Readout is expressed in relative luminescence units (RLU).

**Table 2.** Details of reference antisera obtained from OIE, NIBSC and APHA for strains of IAV (H1-16) and IBV (B/Yam and B/Vic).

| HA Subtype | Antiserum Strain | Source |
| --- | --- | --- |
| H1 | A/duck/Italy/447/2005 (H1) | OIE |
| H2 | A/duck/Germany/1215/1973 (H2) | OIE |
| H3 | A/psittacine/Italy/2873/2000 (H3) | OIE |
| H4 | A/cockatoo/England/1972 (H4) | OIE |
| H5 | A/chicken/Scotland/1959 (H5) | APHA |
| H6 | A/turkey/Canada/1965 (H6) | OIE |
| H7 | A/Anhui/1/2013 (H7) | NIBSC |
| H8 | A/turkey/Ontario/6118/1968 (H8) | OIE |
| H9 | A/mallard/Italy/3817-34/2005 (H9) | OIE |
| H10 | A/ostrich/South Africa/2001 (H10) | OIE |
| H11 | A/duck/Memphis/546/1974 (H11) | OIE |
| H12 | A/duck/Alberta/60/1976 (H12) | OIE |
| H13 | A/gull/Maryland/704/1977 (H13) | OIE |
| H14 | A/mallard/Gurjev/263/1982 (H14) | OIE |
| H15 | A/shearwater/Australia/2576/1979 (H15) | OIE |
| H16 | A/gull/Denmark/68110/2002 (H16) | OIE |
| H17 | Polyclonal sera (BATS) | APHA |
| B/YAM | B/Phuket/3073/2013 | NIBSC |
| B/VIC | B/Brisbane/60/2008 | NIBSC |

**Table 3.** Titers in relative luminescence units/mL (RLU/mL) of IAV and IBV hemagglutinin pseudotyped viruses as indicated in **Figure 3**. Protease utilized to achieve the highest titers is indicated. TMPRSS4 is abbreviated to T4 and TMPRSS2 to T2.

| Group I IAV HA |  |  | Group II IAV HA |  |  |
| --- | --- | --- | --- | --- | --- |
| HA Envelope | Titer (RLU/mL) | Protease | HA Envelope | Titer (RLU/mL) | Protease |
| H1 | 2.25E+08 | T4 | H3 | 5.39E+10 | T2 |
| H2 | 6.62E+07 | T2 | H4 | 6.62E+07 | T2 |
| H5 | 1.32E+09 | * | H7 | 5.25E+10 | * |
| H6 | 2.35E+10 | T4 | H6 | 2.35E+10 | T4 |
| H8 | 4.75E+10 | T4 | H10 | 2.68E+10 | T4 |
| H9 | 4.88E+08 | T4 | H14 | 2.92E+10 | T4 |
| H11 | 8.78E+09 | T4 | H15 | 5.16E+10 | T4 |
| H12 | 1.21E+10 | T4 | IBV HA |  |  |
| H13 | 1.44E+09 | T4 | B pre-split | 3.87E+10 | HAT |
| H16 | 5.81E+09 | T4 | B/Vlc-like | 2.89E+10 | HAT |
| H17 | 2.94E+08 | HAT | B/Yam-like | 1.78E+09 | HAT |
| H18 | 5.33E+07 | T4 |  |  |  |

(\*) indicates highly pathogenic avian influenza (HPAI) strains which did not require protease for production

If production titers were less than 5⨯10^7^ RLU/mL, we additionally tested with Transmembrane Serine Protease 2 (TMPRSS2) for the PV strain in the same plasmid ratios (**Figure 2b**). Generally, TMPRSS4 produced the highest titers (RLU/mL) for all subtypes except for H17 and IBV lineages, where HAT produced the highest titers and H2, H3, and H4 required TMPRSS2 [73] for optimal production (**Table 3**). Protease was not necessary for production of HPAI representative viruses, H5 and H7 in **Figure 3** and **Table 2** (as indicated by *). Optimized conditions were then recorded, and PV production volume was scaled up to produce larger PV stocks.

Our optimized method enabled us to produce the most comprehensive pseudotype library to date with representative strains from IAV subtypes H1-H18 and both IBV lineages. **Figure 4** illustrates the range of IAV subtypes already present and available in this library. Full details of current library at the VPU are indicated in **Supplementary Table 1**. These include low pathogenic avian influenza (LPAI) strains from H5 and H7, in addition to HPAI H5 and H7 presented (**Figure 3, Table 3**).

**Figure 4.**
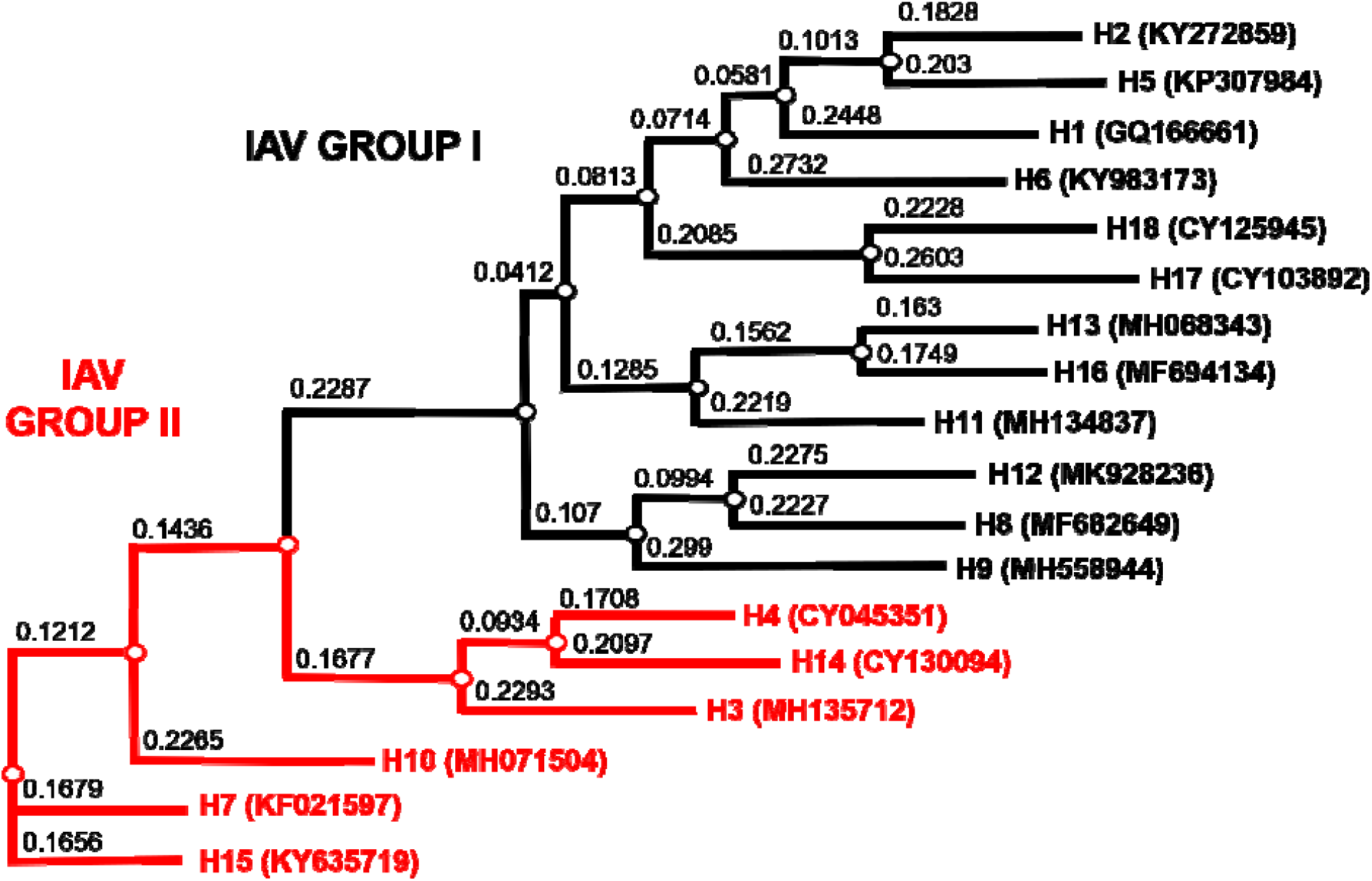
Phylogenetic tree of representative IAV HA from the PV library constructed as shown in **Figure 3** and **Table 3**. Influenza A Group I HA PV are shown in black, IAV Group II PV in red. Accession numbers are reported with the subtype on the tree tips. Nodes are shown at the ends of branches which represent sequences or hypothetical sequences at various points in evolutionary history. Branch lengths indicate the extent of genetic change. The tree generated was constructed with PhyML on the Influenza Research Database (IRD) [69] and graphically elaborated with Archaeopteryx.js (https://sites.google.com/site/cmzmasek/home/software/archaeopteryx-js).

### 3.2. Neutralization of pseudotypes by reference antisera

The neutralization susceptibility of representative PV generated to available HA subtype specific reference antisera (**Table 2**) was assessed. All reference antisera were able to neutralize subtype homologous PV they were tested against (**Figure 5**). We have shown neutralization dose response curves for PV representing IAV strains which have been reported as the cause of human disease, including avian subtypes that have caused zoonotic infection without being associated with sustained human to human transmission (HPAI H5 and H7, and H9) (**Figure 5a**). We have also tested against HA PV that have been associated with swine and human infection, H1 strains which have been found in pigs, which may acquire the ability to transmit to humans due to possible antigenic shift (**Figure 5b**), and avian IAV subtypes which are found in their natural reservoir, wild, and occasionally domesticated, birds (**Figure 5c**) and may evolve in future to novel pandemic strains in humans. Currently there is no commercially available H17 or H18 subtype antisera. Due to the association of H17 in frugivorous bat species, sera collected from bats in Nigeria, as provided by the Animal and Plant Health Agency (APHA), were assessed against the H17 pseudotype (**Figure 5d**). Three bats within a larger panel (only 5 samples shown here) neutralized the H17 PV (**Figure 5d**). Human IBV PV from both Yamagata and Victoria-like lineages were also susceptible to neutralization from reference antisera (**Figure 5e**). Antisera to pre-split IBV strains were not available for this study, nonetheless, our representative pre-split IBV (B/Hong Kong/73) strain was neutralized by both anti-Yamagata and anti-Victoria reference sera. However, this PV was more susceptible to neutralization by antisera generated using HA from the Victoria-like lineage than the Yamagata-like lineage (**Figure 5e**), with IC_50_ dilution values of 688.8 and >10,000, respectively.

**Figure 5.**
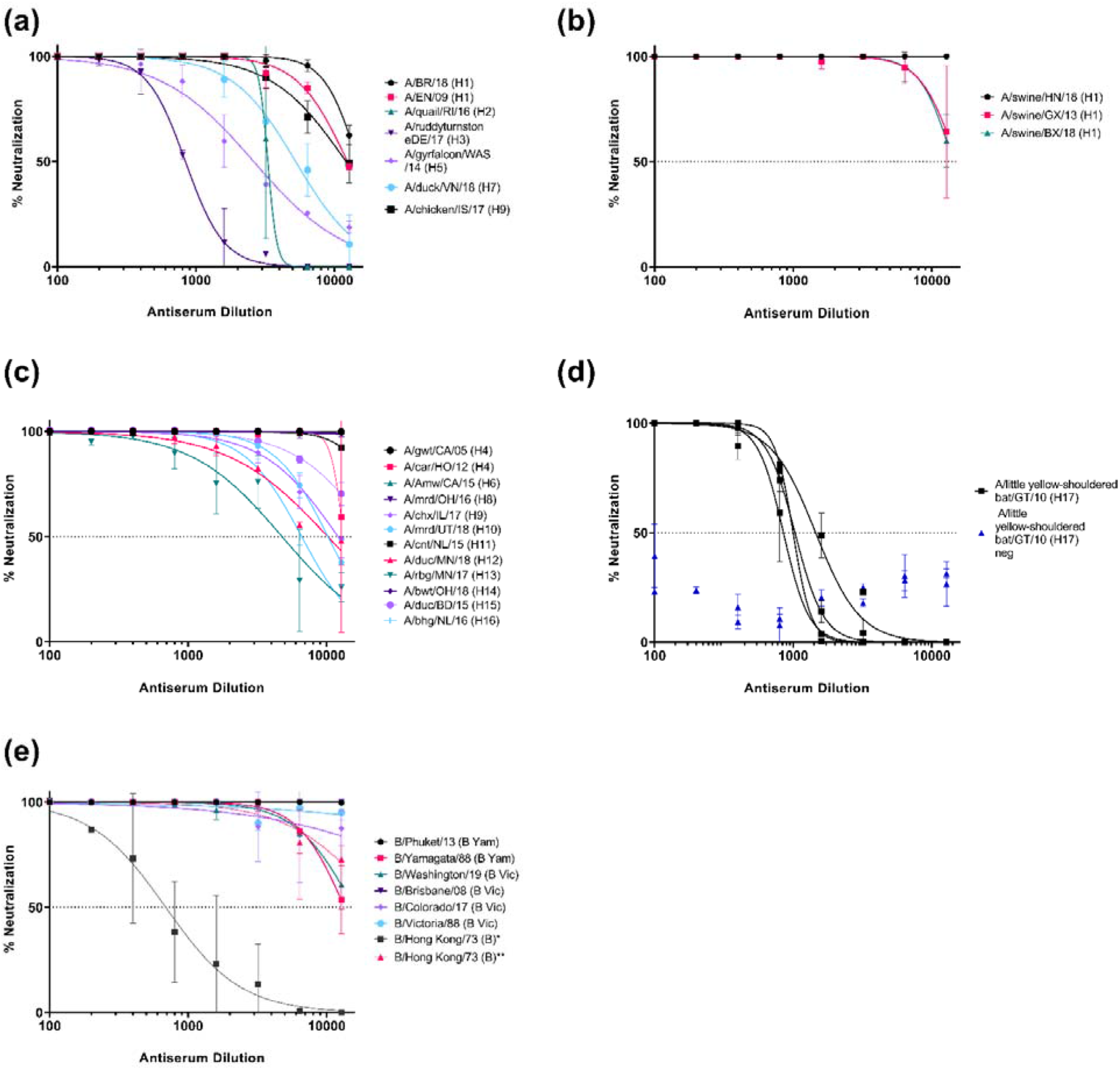
Neutralization of influenza pseudotypes by reference antisera and bat sera from influenza surveillance. (**a**) Neutralization of representative IAV subtypes which have previously caused infection in humans (H1, H2, H3, H5, H7, and H9). (**b**) Neutralization of pseudotypes representing IAV isolated from swine (H1). (**c**) Neutralization of pseudotypes which are representative of IAV found in avian populations (H4, H6, H8, H9, H10, H11, H12, H13, H14, H15 and H16). (**d**) Neutralization of H17 PV (A/little yellow-shouldered bat/Guatemala/060/2010) by bat sera from bat surveillance sampling in Nigeria as provided by APHA. (**e**) Neutralization of IBV pseudotypes which have caused human infection (B/Yamagata-like and B/Victoria-like viruses and pre-split IBV). As pre-split antiserum was not available, neutralization susceptibility of this PV to B/Yamagata lineage antisera (*) and B/Victoria lineage antisera (**) have been shown. Neutralization was measured by a luciferase reporter assay. Reference antisera and bat sera were serially diluted two-fold from a starting dilution of 1:100. 1.0×10^6^ RLU of PV was then added to each well. For all plots, each point represents the mean and standard deviation of two replicates per dilution. Details of reference antisera are indicated in **Table 2**.

### 3.3. Mouse immunogenicity studies

We have conducted preliminary mouse immunogenicity studies to determine the capacity of potential selected HA vaccine candidates to elicit a measurable immune response, assess safety, and to see if our dose and dosing regimen could inform future pre-clinical trials for protection and efficacy. In this study, we have measured immune responses to vaccination in mouse sera (humoral immune responses). The humoral immune response is assessed from the post-vaccination appearance of antibody directed at the specific vaccine antigen at appointed time points. Using our PV library, we have measured functional antibodies in mouse sera that can be applied to many samples in low containment, which would not be possible using wildtype viruses.

We first determined if our dose of 50 µg pEVAC HA DNA could generate an immune response that could be monitored across a certain time frame, and if additional immunizations could increase this specific immune response. It should be noted that all mice used for our experiments were naïve and have not had prior exposure to influenza, hence it is assumed that neutralization of PV will be due to the immune response generated by immunization with influenza antigens.

We observed an immune response in all mice (n=6) vaccinated with pEVAC EN/09 (H1) against the corresponding homologous EN/09 (H1) PV in all post-vaccination samples from the earliest time of sampling (42 dpi) as compared to pre-vaccinated sera (day 0) (**Figure 6**). At 42 dpi, mice had received 3 immunizations (day 0, day 14, and day 28) and had already developed neutralizing antibody responses compared to day 0 (**Figure 6a**). At 56 and 70 dpi, mice had received 4 immunizations (day 0, day 14, day 28, and day 42) (**Figure 6b-c**). A significant increase in detectable neutralizing antibodies can be seen between the third and fourth round of immunizations (**Figure 6d**) demonstrating that boosting the immune response with subsequent immunizations may give rise to stronger neutralizing titers. There was no significant difference between neutralizing activity of mouse sera from bleeds taken at 56 and 70 dpi (**Figure 6d**), suggesting that we had employed an ideal number and interval of immunizations of pEVAC HA to achieve optimal strain-specific titers in mice.

**Figure 6.**
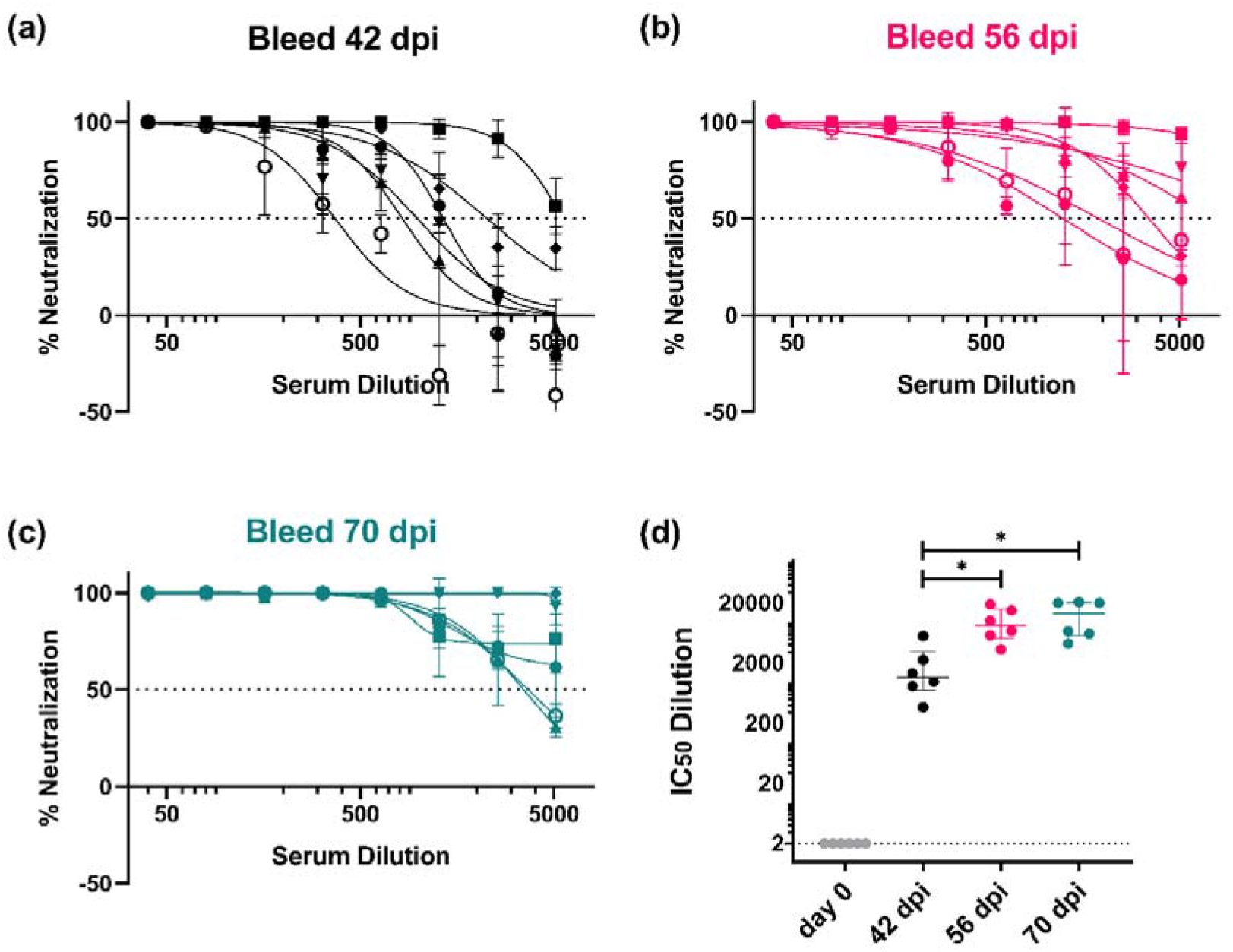
*In vitro* neutralizing activity of mouse sera against A/England/195/09 (H1) (EN/09) as monitored at specific timepoints during the immunization protocol. Mice were vaccinated with 50 µg pEVAC – EN/09 (H1) on days 0, 14, 28, and 42. Bleeds were taken (**a**) 42 days post immunization (dpi), (**b**) 56 dpi, and (**c**) 70 dpi (terminal bleed). Neutralizing activity was tested against 1×10^6^ RLU of A/England/195/09 (H1) PV. (**d**) Comparison of half maximal inhibitory dilutions (IC_50_) in post-vaccination samples as a function of time is shown in brackets (*p < 0.05). Broken line shows an assigned baseline level of 2 indicating 0% neutralization. For plots (**a-c**), the mean and standard deviation of individual mouse serum samples are shown (n=6). Plot (**d**) shows the median and interquartile range of samples tested.

The ability of mouse sera vaccinated with a specific IAV strain to neutralize strains of the same subtype was then evaluated. This was to assess possible strain cross-reactivity of the immune responses elicited by vaccination to these PV. This is especially important for influenza, which is subject to continuous random antigenic drift and wherein viruses of the same subtype may belong to different clades. Given this, we chose to investigate H1, the most recent IAV pandemic strain (in 2009), and HPAI subtypes H5 and H7, which have caused human spillovers from fatal poultry outbreaks in the past. We employed the same immunization procedure as detailed above and terminal bleeds were assessed for neutralizing activity.

For our A/H1 panel, we immunized mice with the pandemic strain EN/09 (H1) (**Figure 7a**). We tested against a previous H1 pandemic strain, SC/1918 (Spanish flu) and a possible emerging pandemic strain, swine/BJ/18, guided by the knowledge that the last H1N1 pandemic (swine flu) was caused by a quadruple-reassortant virus, containing genes from Asian and European swine, North American avian as well as human influenza virus [30]. Terminal sera from mice immunized with EN/09 were able to neutralize all H1 PV tested, with no significant difference observed in neutralization activity against homologous and heterologous strains of the same subtype, with representative PV strains covering 100 years, from 1918 to 2018 (**Figure 7a**).

**Figure 7.**
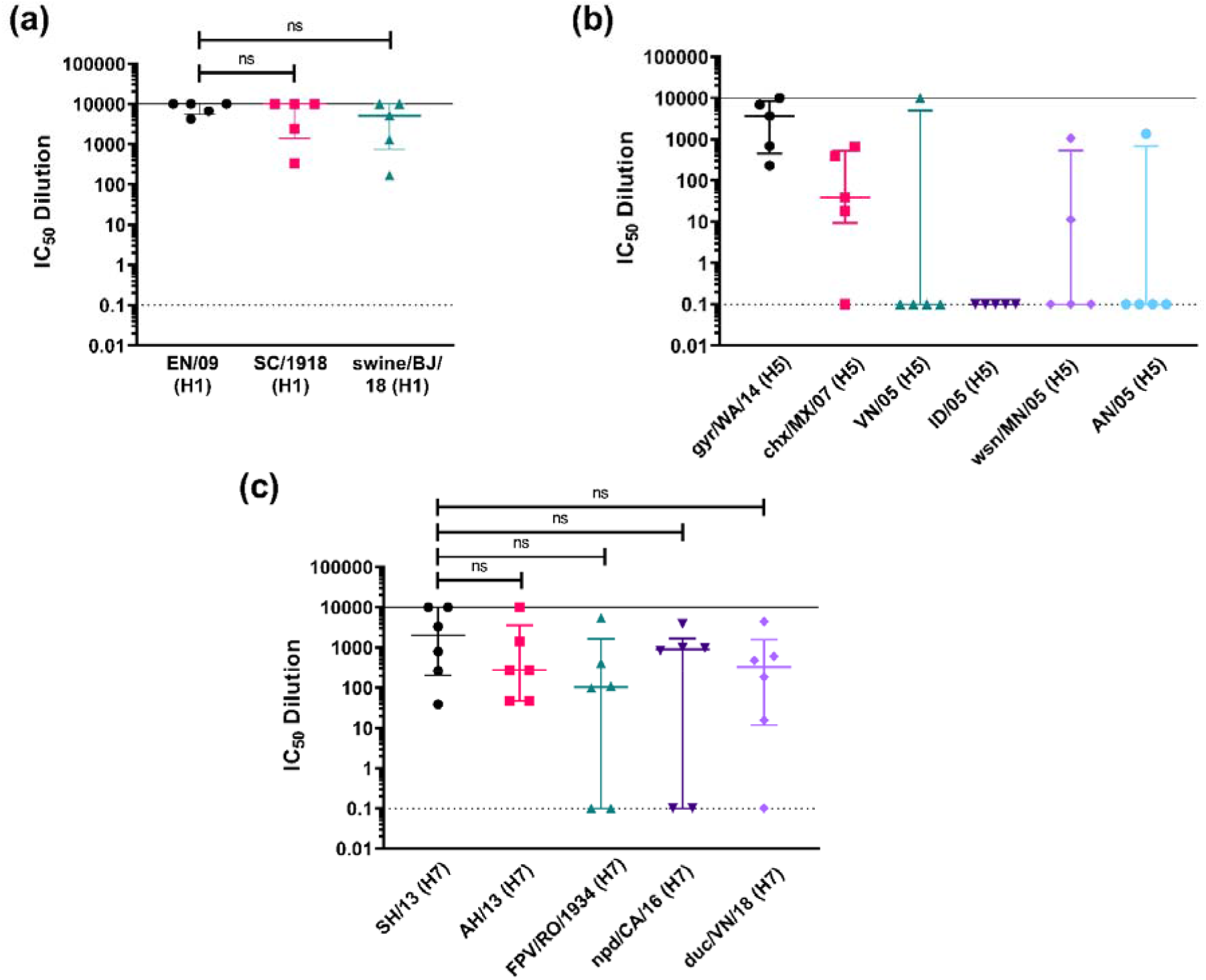
*In vitro* neutralizing activity as shown by IC_50_ dilution of mouse sera vaccinated with an HA subtype tested against homologous PV and representative PV strains of the same subtype. (**a**) Mice were vaccinated with 50 µg pEVAC HA A/England/195/09 (H1) (EN/09) (n=5). Terminal bleeds (70 dpi) were tested against H1 PV strains (x-axis): homologous EN/09, A/South Carolina/1/1918 (H1) (SC/1918) and A/swine/Beijing/301/18 (H1) (swine/BJ/18). (**b**) Mice were vaccinated with 50 µg pEVAC HA A/gyrfalcon/Washington/41088-6/14 (H5) (gyr/WA/14) (n=5). Terminal bleeds (70 dpi) were tested against H5 PV strains (x-axis): homologous gyr/WA/14, A/chicken/Mexico/7/07 (H5) (chx/MX/07), A/Indonesia/5/05 (H5) (ID/05), A/Vietnam/1203/04 (H5) (VN/05), A/whooper swan/Mongolia/244/05 (H5) (wsn/MN/05), and A/Anhui/1/05 (H5) (AN/05). (**c**) Mice were vaccinated with 50 µg pEVAC HA A/Shanghai/2/13 (H7) (SH/13) (n=6). Terminal bleeds (70 dpi) were tested against H7 PV strains (x-axis): homologous SH/13, A/Anhui/1/13 (H7) (AH/13), A/FPV/Rostock/1934 (H7) (FPV/RO/1934), A/northern pintail duck/California/UCD1582/16 (H7) (npd/CA/16), and A/duck/Vietnam/HU10-64/18 (H7) (duc/VN/18). For all plots, the median and interquartile range of individual mouse serum samples per immunization group are shown. Solid line indicates an assigned maximum IC_50_ dilution of 10,000 showing 100% neutralization and broken line shows an assigned baseline level of 0.1 indicating 0% neutralization (cell only mean). Comparisons of no significant difference (ns: p>0.05) against the homologous PV are shown in brackets.

We immunized mice with gyr/WA/15 (H5) for our A/H5 panel (**Figure 7b**). We tested across six different clades, gyr/WA/14 (clade 2.3.4.4c), chx/MX/07 (American non-goose Guangdong), ID/05 (clade 2.1.3.2), VN/04 (clade 1), wsn/MN/05 (clade 2.2), and AN/05 (clade 2.3.4). HPAI H5 strains have been known to cause deadly outbreaks in poultry with some human spillover in the past [74, 75]. H5 viruses especially those in clade 2 are known to evolve rapidly and extensively, with newly emerging strains circulating in many regions of the world [74]. Our findings here demonstrate that terminal sera from mice immunized with gyr/WA/15 (H5) were unable to neutralize the other H5 PV tested as effectively as the homologous strain used for vaccination (**Figure 7b**). Interestingly, one mouse developed a broadly neutralizing response and was able to neutralize all PV tested except for IN/05 but all other samples revealed no H5 cross-strain neutralizing immune response (**Figure 7b**).

Mice were immunized with SH/13 (H7) for the H7 panel (**Figure 7c**). In addition to the homologous SH/13 (H7) PV, we tested against four other H7 strains, FPV/RO/1934, the historical H7 fowl plague virus of 1934, a human IAV PV, AH/13, and two avian PV, npd/CA/16, and duc/VN/18. Terminal sera from mice immunized with SH/13 were able to neutralize all H7 PV tested, with no significant difference observed in the means of the IC_50_ dilution values obtained against homologous and heterologous strains of the same subtype (**Figure 7c**). Some serum samples were unable to neutralize all three H7 avian PV, but all serum terminal bleeds were effective against the other H7 human PV tested, AH/13, with neutralization of the homologous strain showing the same pattern (**Figure 7c**).

We then examined the breadth of responses within a subtype with the idea that one vaccination could protect for small changes caused by antigenic drift as well as providing some initial protection from reassortant viruses which can transmit between species. This is also the basis of strain selection for seasonal influenza vaccination [54, 76]. To test this, we examined cross strain neutralization in mice vaccinated with antigens from strains of IAV H3 isolated from human and avian origins. H3 circulates in the human population and is a component of the quadrivalent influenza vaccine, transmission is often from animal sources [6, 7], and therefore cross-reactive immune responses would be beneficial.

We immunized groups of mice with H3 from avian strains, A/ruddy turnstone/ Delaware Bay/606/2017 (H3) (rtn/DE/17) and A/duck/Quang Ninh/220/2014 (H3) (duc/QN/14), and 2 strains of H3 of clades 3C.2a2 which have circulated recently in the human population, A/South Australia/34/2019 (H3) (SA/19) and A/Switzerland/8060/2017 (H3) (SZ/17), respectively (**Figure 8**). Terminal sera from mice were tested against two avian H3 PV matched to the immunization antigens, rtn/DE/17 and duc/QN/14, and one representative human strain PV, A/Udorn/307/1972 (H3).

**Figure 8.**
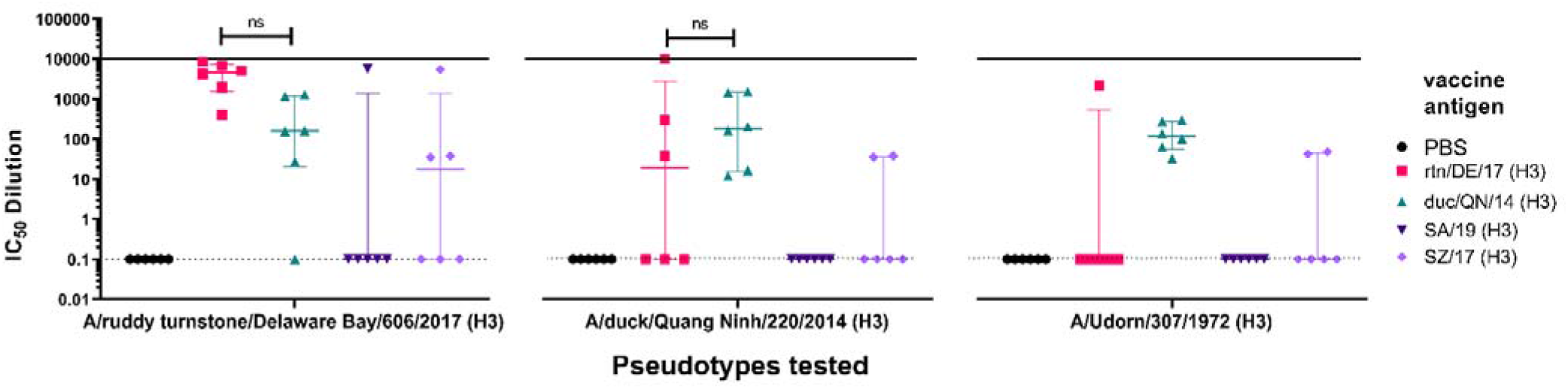
*In vitro* neutralizing activity as shown by IC_50_ dilution of mouse sera vaccinated with avian and human pEVAC H3 vaccine antigens tested against homologous avian H3 PV and a representative human PV strain of H3. Four groups consisting of 6 mice each (n=6/group) were vaccinated with 50 µg pEVAC HA-A/ruddy turnstone/Delaware Bay/606/2017 (H3) (rtn/DE/17) (pink square) or HA-A/duck/Quang Ninh/220/2014 (H3) (duc/QN/14) (green triangle), HA-A/South Australia/34/2019 (H3) (SA/19) (violet inverted triangle), and HA-A/Switzerland/8060/2017 (H3) (SZ/17) (purple diamond), respectively. An additional group of mice was vaccinated with PBS (negative control group) (n=6). Terminal bleeds (70 dpi) were tested against H3 PV strains: 2 homologous avian PV, rtn/DE/17 and duc/QN/14, and one human PV, A/Udorn/307/1972 (UD/1972), as shown in the x axes. For all plots, the median and interquartile range of individual mouse serum samples per immunization group are shown. Solid line indicates an assigned maximum IC_50_ dilution of 10,000 showing 100% neutralization and broken line shows an assigned baseline level of 0.1 indicating 0% neutralization (cell only mean). Comparisons of no significant difference (ns: p>0.05) against the homologous PV are shown in brackets.

Terminal sera from mice immunized with rtn/DE/17 (**Figure 8, 1**^**st**^ **panel**) were able to strongly neutralize homologous PV (rtn/DE/17) with an IC_50_ dilution range of 413-8634. These mice were also able to neutralize heterologous PV duc/QN/14 with no significant difference compared to sera from mice vaccinated with duc/QN/14 antigen (**Figure 8, 2**^**nd**^ **panel**). Only one mouse from the group vaccinated with rtn/DE/17 produced responses which were able to neutralize human H3 PV, A/Udorn/307/1972 (**Figure 8, 3**^**rd**^ **panel**), whereas, all mice vaccinated with duc/QN/14 were able to neutralize A/Udorn/307/1972. This is promising as vaccination with an avian H3 has shown neutralization of a human H3 PV, albeit an older strain from 1972. Mice immunized with SA/19, a human H3 antigen, did not neutralize any of the PV tested except for serum from one mouse which was able to neutralize PV rtn/DE/17 (IC_50_ dilution ∼5659). Sera from half of the mice immunized with SZ/17 were able to neutralize PV rtn/DE/17 and 2 mice were able to neutralize duc/QN/14 and PV A/Udorn/307/1972. These results suggest that vaccination with duc/QN/14 showed the best immune responses against all H3 PV tested, either avian or human (**Figure 8**).

As seen with results indicated above (**Figures 7** and **8**), vaccination with subtype specific antigens have very little effect against other strains from that same subtype. This is what is observed in seasonal vaccination that generates subtype-specific antibodies that will have little or no efficacy against drifted strains [77, 78]. An immunization that gives rise to broadly protective humoral immunity against influenza remains a sought-after goal. With this in mind, we have attempted to demonstrate cross-subtype neutralization from immunization among IAV subtypes that are closest to each other on the phylogenetic tree for IAV (**Figure 4**) and between the two IBV lineages, B/Victoria (B/Vic) and B/Yamagata-like viruses (B/Yam).

For the IAV H7/H10/H15 study (**Figure 9a, 1**^**st**^ **panel**), mice vaccinated with npd/CA/16 (H7) showed no significant difference in neutralizing activity (n.s.) with mice vaccinated with duc/VN/18 (H7) when tested against the duc/VN/18 (H7) PV. This was also observed with mice vaccinated with mrd/UT/18 (H10) and duc/BD/15 (H15) (**Figure 9a, 1**^**st**^ **panel**). This suggests that vaccination with northern pintail duck/CA/16 (H7), mrd/UT/18 (H10), and duc/BD/15 (H15) produced a similar neutralizing response to the homologous antigen against the duc/VN/18 (H7) PV *in vitro*. Mice vaccinated with the other antigens duc/BD/15 (H10) and wts/WAUS/79 (H15) showed little to no neutralization of the H7 PV, this may be partly due to this H15 virus being isolated in 1979, suggesting that this avian H15 diverged between 1979 and 2018 (**Figure 9a, 1**^**st**^ **panel**).

**Figure 9.**
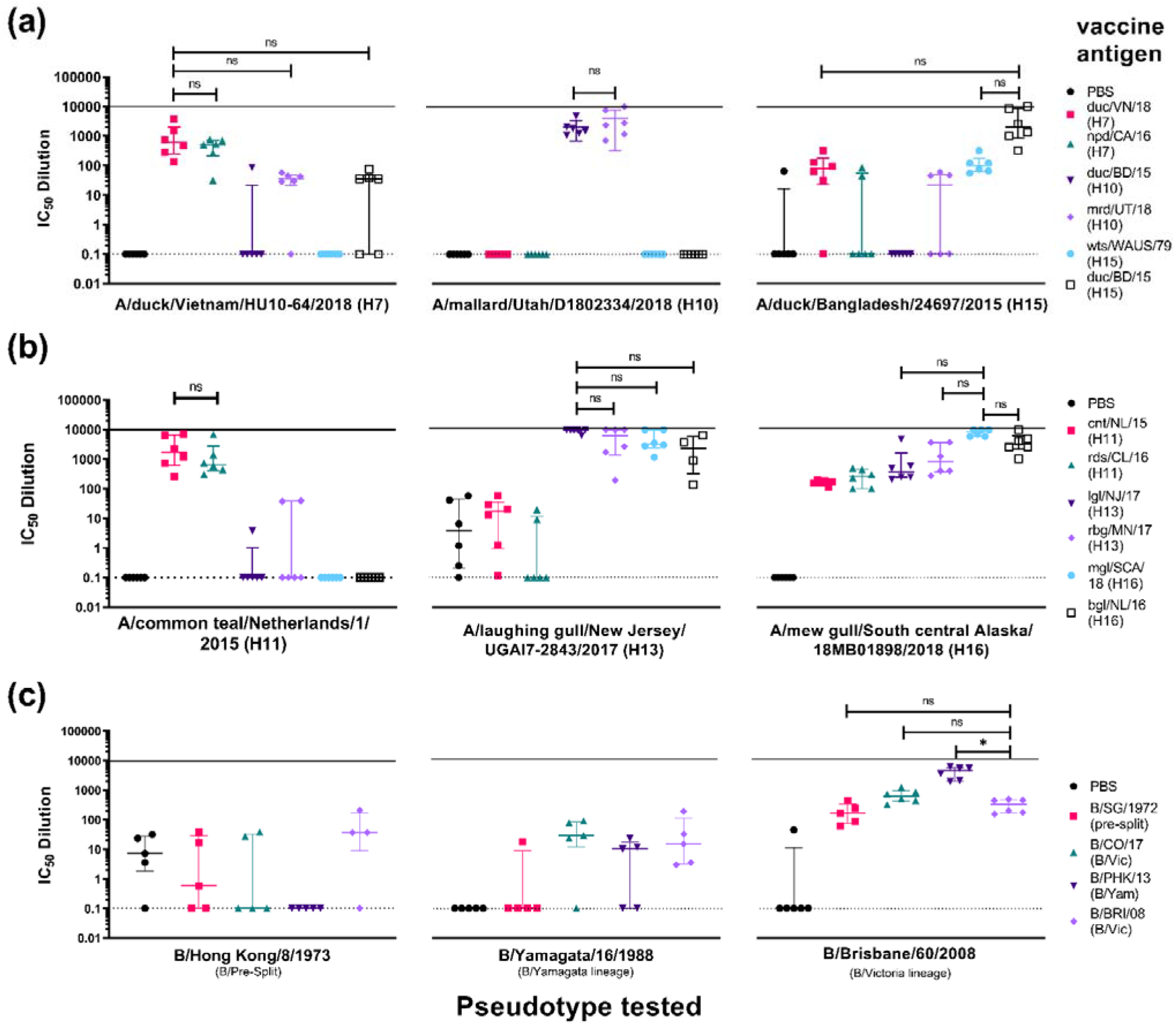
*In vitro* neutralizing activity as shown by IC_50_ dilution of mouse sera vaccinated with HA antigens from closest phylogenetically related IAV subtypes, (**a**) H7, H10, and H15, (**b**) H11, H13 and H16, and (**c**) IBV HA antigens from pre-split, Yamagata and Victoria like-lineages. Neutralizing activity of sera from vaccinated mice were tested against homologous and heterologous strains from the same subtype and a representative strain within the related subtypes. (**a**) IAV H7/H10/H15 study. Six groups consisting of 6 mice each (n=6/group) were vaccinated with 50 µg pEVAC expressing A/duck/Vietnam/HU10-64/2018 (H7) (duc/VN/18) (pink square), A/northern pintail duck/California/UCD1582/2016 (H7) (npd/CA/16) (green triangle) (n=6), A/duck/Bangladesh/24268/2015 (H10) (duc/BD/15) (violet inverted triangle), A/mallard/Utah/D1802334/2018 (H10) (mrd/UT/18) (purple diamond), A/wedge-tailed shearwater/Western Australia/2576/1979 (H15) (wts/WAUS/79) (light blue circle), and A/duck/Bangladesh/24697/2015 (H15) (duc/BD/15) (hollow black square), respectively. Terminal bleeds (70 dpi) were tested against H7 PV duc/VN/18, H10 PV mrd/UT/18, and H15 PV duc/BD/15 as shown on the x-axes. (**b**) IAV H11/H13/H16 study. Six groups consisting of 6 mice each (n=6/group) were vaccinated with 50 µg pEVAC cloned with A/common teal/Netherlands/1/2015 (H11) (cnt/NL/15) (pink square), A/red/shoveler/Chile/C14653/2016 (H11) (rds/CL/16) (green triangle), A/laughing gull/New Jersey/UGAI7-2843/2017 (H13) (lgl/NJ/17) (violet inverted triangle), A/ring-billed gull/Minnesota/OPMNAI0816/2017 (H13) (rbg /MN/17) (purple diamond), A/black-headed gull/Netherlands/1/2016 (H16) (bhg/NL/16) (light blue circle), and A/mew gull/South central Alaska/18MB01898/2018 (H16) (mgl/SCA/18) (black hollow square). Terminal bleeds (70 dpi) were tested against H11 PV cnt/NL/15, H13 PV lgl/NJ/17, and H16 PV mgl/SCA/18 as shown in the x-axes. (**c**) IBV cross lineage study. Four groups of mice were vaccinated with 50 µg pEVAC cloned with B/Singapore/222/1979 (pre-split) (B/SG/1979) (pink square) (n=5), B/Colorado/06/2017 (B/Vic) (B/CO/17) (green triangle) (n=6), B/Phuket/3073/2013 (B/Yam) (B/PHK/13) (violet inverted triangle) (n=6), and B/Brisbane/60/2008 (BVIC) (B/BRI/08) (purple diamond) (n=6), respectively. Terminal bleeds (70 dpi) were tested against a representative B pre-split PV, B/Hong Kong/8/1973, a representative B/Yamagata PV, B/Yamagata/16/1988, and B/Victoria PV B/Brisbane/60/2008 as shown in the x-axes. For all plots, an additional group of mice was vaccinated with PBS (n=5/6). Plots show the median and interquartile range of individual mouse serum samples per immunization group. Solid line indicates an assigned maximum IC_50_ dilution of 10,000 showing 100% neutralization and broken line shows an assigned baseline level of 0.1 indicating 0% neutralization. Comparisons of no significant difference (ns: p>0.05) and significant difference (* p<0.05) among IC_50_ values with antigen homologous to the PV being tested against is shown in brackets. In the case of (**c**), comparison is made with neutralization of antigen belonging to the same lineage as the PV it is tested against.

Results for groups tested against mrd/UT/18 (H10) PV are more clear-cut (**Figure 9a, 2**^**nd**^ **panel**), with only groups vaccinated with H10 antigens showing neutralizing activity against the PV. There is also no significant difference between the IC_50_ values against the mrd/UT/18 (H10) PV in the group vaccinated with the other H10, duc/BD/15, to that vaccinated with the homologous mrd/UT/18 (H10) (**Figure 9a, 2**^**nd**^ **panel**). Here, only neutralization of PV by mice vaccinated with the same subtype is demonstrated. Looking at the phylogenetic tree (**Figure 4**), H10 resides on a different branch than H7 and H15, and therefore it was highly unlikely that cross-subtype neutralization would be observed.

For groups tested against duc/BD/15 (H15) PV, vaccination with the homologous antigen, the other H15 antigen, wts/WAUS/79, and duck/VN/18 (H7) showed neutralizing activity (**Figure 9a, 3**^**rd**^ **panel**). Neutralizing activity of mice vaccinated with all other antigens tested was closer to that of the negative control group (PBS), although a few responders, located above the upper extreme quartile, were observed.

For the IAV H11/H13/H16 study (**Figure 9b, 1**^**st**^ **panel**), mice vaccinated with rds/CL/16 (H11) showed no significant difference in neutralizing activity (n.s.) with mice vaccinated with cnt/NL/15 (H11) when tested against the cnt/NL/15 (H11) PV. This suggests that vaccination with rds/CL/16 (H11) produces the same neutralizing response as its homologous antigen against the cnt/NL/15 (H11) PV *in vitro*. Mice vaccinated with both H13 antigens showed very little neutralization against the H11 PV, with only two mice of the lgl/NJ/17(H13) group and one mouse from the rbg/MN/17 (H13) group showed neutralization though IC_50_ values were closer to the negative control group (**Figure 9b, 1**^**st**^ **panel**). There was no neutralization of the H11 PV by mice vaccinated with H16 antigens, mgl/SCA/18 (H16) and bhg/NL/16 (H16) suggesting that this antigen did not elicit significant responses to epitopes which are common to both the H11 and H16 IAV strains.

When sera were tested against lgl/NJ/17 (H13) (**Figure 9b, 2**^**nd**^ **panel**), mice vaccinated with H11 antigens produced poor neutralizing responses. However, similar to what we observed in the H11 PV neutralization (**Figure 9b, 1**^**st**^ **panel**), there was no significant difference between the IC_50_ values of groups vaccinated with the other H13, rbg/MN/17 (H13), to that vaccinated with the homologous lgl/NJ/17 (H13) against the lglNJ/17 (H13) PV (**Figure 9b, 2**^**nd**^ **panel**). In contrast to previous results (**Figure 9b, 1**^**st**^ **panel**), cross subtype neutralizing activity was observed in sera from mice vaccinated with both H16 antigens showing a strong neutralization response, as there was no significant difference between the IC_50_ values of these groups with those vaccinated with the homologous lgl/NJ/17 (H13) against the lgl/NJ/17 (H13) PV (**Figure 9b, 2**^**nd**^ **panel**). Looking at the phylogenetic tree (**Figure 4**), H11 is farther from H13 and H16, which may explain the lack of cross-subtype neutralization seen here. It is of note here that low level background neutralization was observed in sera of the negative control group against laughing gull/NJ/17 (H13) PV, this was not seen when this group was tested against the H11 or H16 PV.

Interestingly, all vaccination groups neutralized mgl/SCA/18 (H16) PV (**Figure 9b, 3**^**rd**^ **panel**). Despite this, vaccinations with the other H16 antigen (bhg/NL/16 (H16)), and both H13 antigens achieved IC_50_ titers which had no significant difference (n.s.) compared to the homologous mgl/SCA/18 (H16) vaccination against the mgl/SCA/18 (H16) PV (**Figure 9b, 3**^**rd**^ **panel**). Neutralizing responses were also observed in sera of mice immunized with H11 antigens, albeit not as strong as that of the homologous, nonetheless these responses were significantly different from the negative control mice sera (p<0.05) (**Figure 9b, 3**^**rd**^ **panel**).

For the IBV study for groups tested against B/Hong Kong/1973 (B/HK/1973) PV, a pre-split IBV (**Figure 9c, 1**^**st**^ **panel**) (n=5 for all groups), no neutralization was observed in sera from mice regardless of the antigen they had been inoculated with including the pre-split antigen B/Singapore/1972 (B/SG/1972). None of the groups showed responses that were significantly different from sera collected from mice in the negative control group (PBS), which, incidentally, was showing some background against this PV. Most of the mice that were vaccinated with any of the antigens tested failed to reach 50% neutralization against the pre-split PV (**Figure 9c, 1**^**st**^ **panel**).

When groups were tested against B/Yam/1988 (B/Yam) (**Figure 9c, 2**^**nd**^ **panel**) (n=5), sera from mice vaccinated with B/SG/1972 (pre-split) did not show any neutralization, except for one outlier that was outside the upper extreme quartile. Neutralization was observed for sera from all other groups including, as expected, those vaccinated with the antigen from Yamagata-like lineage (B/PHK/13) (**Figure 9b, 2**^**nd**^ **panel**). Nonetheless, vaccination employing these antigens did not produce strong neutralizing responses against the B/Yam PV, as responses showed no significant difference with the PBS group, with IC_50_ dilution values ranging from 0.1-100.

Results for groups tested against B/Bri/08 PV (**Figure 9c, 3**^**rd**^ **panel**) were interesting; as sera from mice in all groups were able to neutralize this B/Vic PV. Sera from mice vaccinated with B/SG/1972 (pre-split) and the other B/Vic antigen, B/CO/17, achieved IC_50_ dilution values which had no significant difference (n.s) compared to that vaccinated with the homologous B/Bri/08 (B/Vic) against the B/Bri/08 (B/Vic) PV (p>0.05). Sera from mice vaccinated with antigens from the other lineage, B/PHK/13 (B/Yam) achieved IC_50_ titers which were higher than those observed for mice from the group vaccinated with homologous B/Bri/08 (BVIC) antigen (*p<0.05) against B/Bri/08 (BVIC) PV. This suggests that vaccination with either pre-split, B/Yam or B/Vic lineage antigens produce a significant neutralizing response against this B/Victoria PV.

### 3.4. In vitro neutralization of HA pseudotypes by HA-stem directed monoclonal antibodies

It is desirable to have antibodies that will elicit a broad, cross-subtype specific response in order to address a pandemic threat. The influenza pseudotype microneutralization (pMN) assay is highly sensitive and specific for detecting neutralizing antibodies against influenza viruses regardless if they are HA-head specific or are targeted against the HA stem, making it an excellent test of antibody functionality *in vitro* [55].

Several broadly reactive monoclonal antibodies have been developed for use in immunotherapy against influenza. Monoclonal antibody CR9114 binds to IBV from both lineages and additionally binds influenza A viruses from both group 1 and group 2 [79], and FI6 is a pan-influenza A neutralizing antibody [80]. Both CR9114 and FI6 bind to a highly-conserved epitope in the HA stem [79, 80] enabling them to broadly neutralize influenza viruses and providing protection against lethal influenza challenge *in vivo*. Here, we show neutralization of representative IAV and IBV PV by both mAbs (**Figure 10, Table 4**).

**Figure 10.**
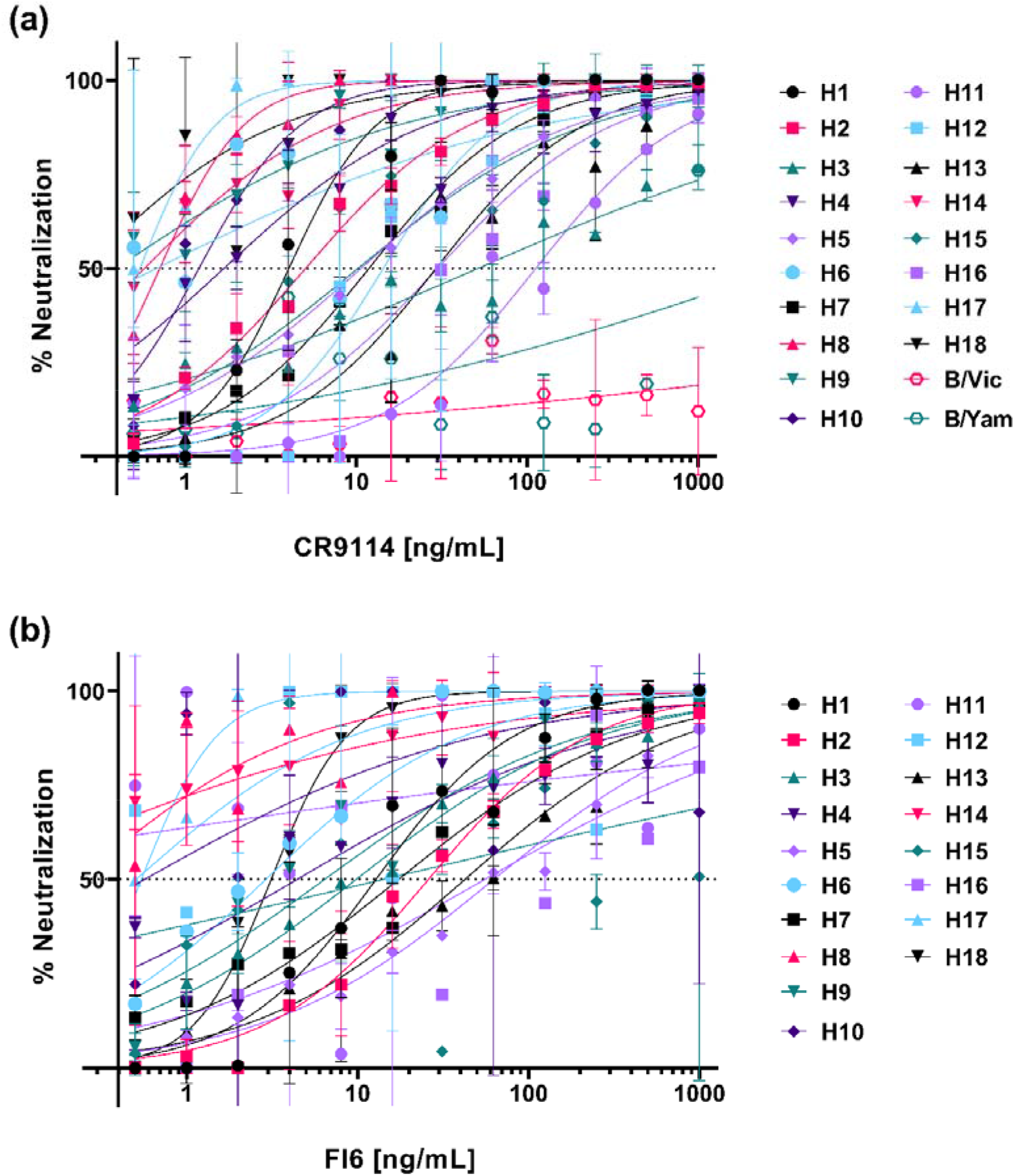
Neutralization of representative IAV and IBV PV *in vitro* by CR9114 and FI6. PV neutralization was measured by a luciferase reporter assay. **(a)** CR9114 and **(b)** FI6 were serially diluted two-fold from a starting concentration of 1000 ng/mL to 0.5 ng/mL against all pseudotypes. An input value of 1.0×10^6^ RLU of PV was then added to each well. For **(a)** and **(b)**, each point represents the mean and standard deviation of two replicates per dilution. IBV strains were not tested against FI6.

**Table 4.** IC_50_ (half-maximal inhibitory concentration) values of CR9114 and FI6 against representative influenza PV *in vitro*. (-) indicates no neutralization. n.d. indicates experiment not done.

| *Pseudotype Virus (PV)* |  | IC<sub>50</sub> [ng/mL] |  |
| --- | --- | --- | --- |
| Subtype | Strain | CR9114 | FI6 |
| H1 | A/England/195/2009 | 3.63 | 13.25 |
| H2 | A/quail/Rhode Island/16-018622-1/2016 | 5.06 | 26.70 |
| H3 | A/ruddy turnstone/Delaware Bay/606/2017 | 51.62 | 9.83 |
| H4 | A/Calidris ruficollis/Hokkaido/12EY0172/2012 | 1.68 | 8.36 |
| H5 | A/gyrfalcon/Washington/41088-6/2014 | 10.74 | 60.15 |
| H6 | A/American wigeon/California/HS007A/2015 | 0.68 | 2.91 |
| H7 | A/Shanghai/02/2013 | 11.88 | 17.39 |
| H8 | A/mallard duck/Ohio/16OS0672/2016 | 0.71 | 0.23 |
| H9 | A/chicken/Israel/291417/2017 | 0.39 | 6.41 |
| H10 | A/mallard/Utah/D1802334/2018 | 1.26 | 0.57 |
| H11 | A/red shoveler/Chile/C14653/2016 | 120.90 | 0.02 |
| H12 | A/duck/Mongolia/850/2018 | 15.06 | 0.51 |
| H13 | A/laughing gull/New Jersey/UGAI17-2843/2017 | 31.97 | 52.05 |
| H14 | A/blue-winged Teal/Ohio/18OS1695/2018 | 0.58 | 0.06 |
| H15 | A/wedge-tailed shearwater/Western Australia/2576/1979 | 10.73 | 14.12 |
| H16 | A/black-headed gull/Netherlands/1/2016 | 45.41 | 55.50 |
| H17 | A/little yellow-shouldered bat/Guatemala/60/2010 | 0.54 | 0.34 |
| H18 | A/flat-faced bat/Peru/030/2010 | 0.26 | 3.14 |
| B | B/Hong Kong/8/1973 | - | n.d. |
| B/Vic | B/Victoria/1/1987 | - | n.d. |
| B/Yam | B/Yamagata/16/1988 | - | n.d. |

Half-maximal inhibitory concentration (IC_50_) of both mAbs against the PV tested were determined. Dose response curves (**Figure 10**) were obtained by normalizing the RLU values against that of the pseudotype only controls corresponding to 0% neutralization and cell-only (no virus) controls corresponding to 100% neutralization. A non-linear regression (curve fit) analysis on the normalized data using a log [inhibitor] versus normalized response variable slope equation to compute for the IC_50_ values was then carried out. The IC_50_ values for CR9114 against all IAV PV tested were in the range of 0.3-120 ng/mL (**Table 4**). The IC_50_ values for FI6 are more varied, with a range of 0.02-60 ng/mL (**Table 4**).

Both CR9114 and FI6 effectively neutralized key Group I Influenza A subtypes, H1, H2, H3, H5, H7, and H9 *in vitro* (**Figure 10, Table 4**). These representative IAV subtypes have been previously detected in the human propulation, including A(H1N1), A(H2N2), and A(H3N2), strains of which have previously caused global pandemics. Both mAbs were also able to neutralize all influenza PV representative strains from known avian subtypes for both IAV Group I (HA6, HA8, H11, H12, H13, and H16) and Group II (H4, H10, H14, and H15) (**Figure 10**). Notably, bat influenza H17 and H18 were also potently neutralized by CR9114 and FI6 (**Table 4**). In contrast, CR9114 and FI6 showed no neutralization activity against any of the influenza B strains tested. This correlates with previous findings of CR9114 being unable to neutralize influenza B viruses *in vitro* as tested using the classic microneutralization and hemagglutination inhibition assays [79]. Some neutralization activity can be seen for CR9114 against B/Phuket/3073/2013, a B/Yamagata-like virus, at the highest concentration tested (1 µg/mL), but there was no dose-response established indicating no true neutralization. As FI6 is only expected to neutralize influenza A viruses, we did not test it against IBV PV.

## 4. Discussion

Influenza infection contributes annually to morbidity and mortality in humans and in wild and domesticated animals worldwide even with vaccination programs already in place. There is additionally the ever-present threat of a pandemic brought about by novel influenza subtypes to which the population has no pre-existing immunity and of which seasonal vaccines may be unable to protect against. Lessons from the past have shown us, that despite our efforts, we are still unprepared to mitigate the devastating loss of life and livelihood when the next influenza pandemic occurs. Protection provided by current seasonal influenza virus vaccines is generally limited and relies on predictive science. Ideally, vaccines should be rapidly generated upon the emergence of a novel threat and should be able to protect against both drifted and shifted strains, and this is the goal of a universal vaccine approach.

To aid in efforts to create a universal influenza vaccine and assist in pandemic preparedness, we have created a comprehensive influenza hemagglutinin pseudotype library. This library enables assessment of responses in lower containment settings thus negating the requirement for BSL3 facilities that are most commonly required when working with high risk influenza subtypes Using pseudotypes also negates the need to isolate live viruses from clinical material, a process that is expensive and can be technically challenging as well as potentially reducing the genetic authenticity of the isolated virus through egg adaptation. Once the HA sequence has been identified, this can be cloned into a suitable plasmid expression vector. Here we have utilized pI.18, pEVAC and phCMV but other plasmids could be employed, and the amount of DNA required determined using the optimization described in this study. This will ensure the rapid production of high quality and high titer PV which can be utilized in a pMN assay following assembly. Addition of a luciferase reporter plasmid produces results that can be determined rapidly using a system which has the potential to be upscaled to high throughput platforms. Additionally, PV can be stored at -80°C for extended periods of time and as was shown with H5, can be lyophilized and stored for up to 4 weeks at 37°C [81]. Lyophilization could expand the potential to investigate and respond to pandemics or other outbreaks from any subtype at speed and without the need for cold chain storage.

The data presented in this study demonstrated the utility, versatility, and ease, of employing influenza hemagglutinin pseudotyped viruses in pre-clinical studies to further vaccine research using reference standards, improve vaccine antigen design, and to evaluate alternative therapies such as that of mAbs, against influenza. We have also shown that PV in this library are suitable for investigation of neutralization of sera collected from different species including mice, bats, sheep and chickens (**Figure 5, Table 2**). The pseudotype library has also been effective for use with neutralizing reference antisera (**Figure 5**), and this is integral to the vaccine strain selection process for seasonal influenza vaccines. Laboratories around the world that are part of the World Health Organization Global Influenza Surveillance and Response System monitor the antigenic phenotypes of circulating viruses to select vaccine strains for upcoming influenza seasons. However, investigation of emerging strains that could cause pandemics is limited, as it is arduous to isolate and propagate wildtype virus to test against. Our influenza pseudotype library can be employed to test protection offered by existing vaccines and antisera used in their selection, in this instance, as tools for surveillance and pandemic preparedness.

We have conducted several preliminary immunogenicity trials with selected vaccine antigens to inform the design of pivotal trials and to provide possible initial evaluation of vaccine efficacy employing our IAV and IBV pseudotypes. Other screening tools employ assays that evaluate the presence of binding antibodies but are unable to determine if these antibodies are functionally useful within samples. A common screening tool is the Enzyme Linked Immunosorbent Assay (ELISA) which can measure total antibody (e.g. total IgG) that binds to selected antigens. However, only a proportion of the total antibody detected will be capable of inhibiting viral infection and this should be heavily taken into consideration when deciding how to measure the humoral immune response. Alternatively, the immune response may be assessed by neutralization assays employing native virus or viral pseudotypes. The latter can be carried out at BSL2 and allow a rapid, reliable, safe and easy assessment of humoral immune responses of vaccine antigens against influenza subtypes which are difficult to isolate and propagate.

We were able to show that immunization with prospective vaccine antigens and subsequent collection of blood serum samples at appropriate time intervals can be used to evaluate immune responses that are relevant to dosing strategies going forward (**Figure 6**). The data generated could inform the appropriate periods between doses and the number of doses that could provide the optimal immune response. In our immune response monitoring study, we found that immunization with 50 µg pEVAC HA four times in 2-week intervals, produced optimal titers after the 4^th^ inoculation with immune responses at 56 and 70 days post immunization showing no significant difference (**Figure 6d**). Additionally, for vaccines, it may also be useful to explore the shortest time frame within which doses may be completed without a detrimental effect on the final immune response. Our results indicate that employing our immunization and dosing strategy, a 4^th^ immunization with pEVAC HA is necessary to achieve maximal titers, as lower titers were achieved with only 3 compared to 4 immunizations (**Figure 6**). This could be extended in the future to explore prime boost regimens with alternative vectors or proteins.

We also ran investigative trials wherein all mice received the same pEVAC HA vaccine antigen and we performed additional testing using relevant representative PV strains belonging to the same subtype (**Figures 7** and **8**). Our findings provide an indication as to whether immunization with a particular strain of the subtype can neutralize drifted strains of the same subtype, which is very important for lasting vaccine efficacy and protection especially in the case of influenza. This additional testing can also provide an assessment of the robustness and breadth of the humoral immune responses elicited by the vaccine to avian and human strains of the same subtype in the case of IAV and can guide vaccine strain or antigen selection in a vaccine to improve or maintain its protective effect.

We have also looked at comparing immune response against phylogenetically related subtypes to investigate cross-subtype neutralization that can be brought about by vaccination (**Figure 9**). Here, we have also selected IAV strains that are not usually studied, H10, H11, H13, H15, and H16, together with IBV from both lineages. Findings may aid in the development of a vaccine for pandemic purposes or inform possible pre-existing immune responses in the population. Immune responses generated by vaccination with an HA antigen against the homologous PV was successful in all groups tested (**Figures 6-9**). These can be used as control groups for future vaccination experiments. We have found that cross-subtype protection is rare and neutralizing responses not as strong as the homologous antigen against the PV. Several vaccinations, that of an H7 antigen which showed neutralization against an H10 and H15 PV (**Figure 9a, 1**^**st**^ **panel**), H13 and H16 antigens showing cross neutralization with each other (**Figure 9b**), and and B/Yamagata-lineage vaccinated mice neutralizing B/Victoria-lineage PV (**Figure 9c**). Nonetheless, our pseudotype library has enabled us to test immune responses brought about by vaccination against a variety of IAV and IBV pseudotypes. This is a promising *in vitro* screening procedure to guide pre-clinical studies.

In addition to vaccination and antiviral drugs [48], the use of recombinant monoclonal antibodies that are broadly neutralizing against influenza is a promising strategy to counter annual epidemics and pandemic threats especially in individuals with severe disease. These mAbs, several of which are already in clinical development, bind to functionally conserved epitopes such as those in the influenza hemagglutinin (HA) stem, thereby providing strain independent protection [79, 80, 82-84]. For a time, discovery of these broadly neutralizing antibodies has been hampered by the lack of assays to properly show neutralization afforded by these mAbs that is exclusive from hemagglutination inhibition of HA-head directed antibodies. Antibodies that target the HA stem do not inhibit hemagglutination inhibition [85] and are thought to neutralize influenza via other mechanisms. We have successfully employed our pseudotype library to investigate neutralizing activity of HA stem-directed mAbs CR9114 and FI6 against representative IAV and IBV PV available. For instance, we could confirm that CR9114, despite binding to IBV HA, does not neutralize IBV *in vitro*. This is an invaluable tool to test functionality of new immunotherapies against influenza *in vitro*.

This library is expanding as influenza continues to undergo antigenic changes to the HA protein. We believe it can be part of a toolbox of assays that can be made available to researchers and will be especially helpful for studies investigating alternative and innovative influenza vaccine targets. This method employs a system that has the potential to be high throughput, can be easily adapted to other reporters such as GFP, and may be incorporated into large scale clinical trials and surveillance programs. The PV can also be further developed in an ELISA where it will display the HA trimer in its native form and can be used to distinguish HA stalk responses and quaternary epitopes. Lentiviral PV can be constructed to display other potential vaccine targets such as NA and in future, we plan to complete a full NA PV library to complement our HA library. Additionally, these PV could be used to observe glycosylation patterns and their influence on neutralization of influenza. We believe that our influenza PV library will be an invaluable tool for research and would be impactful in the development of solutions against the changing face of influenza.

## Supplementary Materials

Supplementary Table S1. List of influenza hemagglutinin pseudotypes (PV) available at the Viral Pseudotype Unit, University of Kent. This will be dynamically updated and latest version posted to figshare: https://figshare.com/authors/Nigel_Temperton/438525

## Author Contributions

Conceived and designed experiments – JMD, KDC, FF, GC, JH, NT. Performed experiments JMD, KDC, GC, MF. Analyzed the data – JMD, KDC, GC, NT. Plasmid preparation – BA, RW. Flu antigen design DW, BA, RW. Reagent provision – RK, JH, AB, CS, VA. Wrote the paper – JMD, KD. Revised the paper – BA, SS, RW, GC, JH, NT. All authors reviewed and accepted the final version of the manuscript.

## Funding

NT: KdC and GC receive funding from the Bill and Melinda Gates Foundation: Grand Challenges Universal Influenza Vaccines Award: Ref: G101404. NT and JMD receive funding from Innovate UK, UK Research and Innovation (UKRI) for the project: Digital Immune Optimized and Selected Pan-Influenza Vaccine Antigens (DIOS-PIVa) Award Ref: 105078. RW receives funding from EC FETopen (Virofight, Grant 899619). CS receives funding from the Department of Science and Technology of South Africa cost Centre P10000029.

## Institutional Review Board Statement

Ethical approval for collection of bat surveillance sera was received from the Animal Ethics Committee and Research Ethics Committee of University of Pretoria, with certificate numbers V092-18 and REC097-18, respectively. Approval to sample bat populations (VDS/194/S.4/11/T/85) was obtained from the Director/Chief Veterinary Officer of Nigeria, Department of Veterinary and Pest Control Services, Federal Ministry of Agriculture and Pest Control Services, Abuja Nigeria. Mouse studies were approved by the Animal Welfare Ethical Review Body, University of Cambridge (Project license P8143424B).

## Conflicts of Interest

The authors declare no conflict of interest. The funders had no role in the design of the study; in the collection, analyses, or interpretation of data; in the writing of the manuscript, or in the decision to publish the results.

## Notes

### Competing Interest Statement

The authors have declared no competing interest.

## References

1. Bouvier, N.M. and Lowen, A.C. Animal Models for Influenza Virus Pathogenesis and Transmission. Viruses, 2010. 2(8): p. 1530–1563.

2. Bouvier, N.M. and Palese, P. The biology of influenza viruses. Vaccine, 2008. 26: p. D49–D53.

3. Cauldwell, A.V., Long, J.S., Moncorgé, O., and Barclay, W.S. Viral determinants of influenza A virus host range. Journal of General Virology, 2014. 95(6): p. 1193–1210.

4. Long, J.S., Mistry, B., Haslam, S.M., and Barclay, W.S. Host and viral determinants of influenza A virus species specificity. Nature Reviews Microbiology, 2019. 17(2): p. 67–81.

5. Tong, S., Zhu, X., Li, Y., Shi, M., Zhang, J., Bourgeois, M., Yang, H., Chen, X., Recuenco, S., Gomez, J., Chen, L.M., Johnson, A., Tao, Y., Dreyfus, C., Yu, W., McBride, R., Carney, P.J., Gilbert, A.T., Chang, J., Guo, Z., Davis, C.T., Paulson, J.C., Stevens, J., Rupprecht, C.E., Holmes, E.C., Wilson, I.A., and Donis, R.O. New world bats harbor diverse influenza A viruses. PLoS Pathog, 2013. 9(10): p. e1003657.

6. World Health Organization. Influenza virus infection in humans. 2014; Available from: http://www.who.int/influenza/human_animal_interface/virology_laboratories_and_vaccines/influenza_virus_infections_humans_feb14.pdf?ua=1.

7. World Health Organization. Influenza (Seasonal). 2018; Available from: http://www.who.int/news-room/fact-sheets/detail/influenza-(seasonal).

8. Hobbelen, P.H.F., Elbers, A.R.W., Werkman, M., Koch, G., Velkers, F.C., Stegeman, A., and Hagenaars, T.J. Estimating the introduction time of highly pathogenic avian influenza into poultry flocks. Sci Rep, 2020. 10(1): p. 12388.

9. Russell, C.J., Hu, M., and Okda, F.A. Influenza Hemagglutinin Protein Stability, Activation, and Pandemic Risk. Trends Microbiol, 2018.

10. Bodewes, R., Morick, D., de Mutsert, G., Osinga, N., Bestebroer, T., van der Vliet, S., Smits, S.L., Kuiken, T., Rimmelzwaan, G.F., Fouchier, R.A.M., and Osterhaus, A.D.M.E. Recurring influenza B virus infections in seals. Emerging infectious diseases, 2013. 19(3): p. 511–512.

11. Osterhaus, A.D., Rimmelzwaan, G.F., Martina, B.E., Bestebroer, T.M., and Fouchier, R.A.M. Influenza B virus in seals. Science (New York, N.Y.), 2000. 288(5468): p. 1051--1053.

12. de Vries, R.P., de Vries, E., Bosch, B.J., de Groot, R.J., Rottier, P.J.M., and de Haan, C.A.M. The influenza A virus hemagglutinin glycosylation state affects receptor-binding specificity. Virology, 2010. 403(1): p. 17–25.

13. Skehel, J.J. and Wiley, D.C. Receptor Binding and Membrane Fusion in Virus Entry: The Influenza Hemagglutinin. Annu. Rev. Biochem., 2000. 69: p. 531–69.

14. Wiley, D.C. and Skehel, J.J. The Structure and Function of the Hemagglutinin Membrane Glycoprotein of Influenza Virus. Ann. Rev. Biochem, 1987. 56: p. 365–94.

15. Gamblin, S.J. and Skehel, J.J. Influenza hemagglutinin and neuraminidase membrane glycoproteins. J Biol Chem, 2010. 285(37): p. 28403–9.

16. Russell, R.J., Gamblin, S.J., Haire, L.F., Stevens, D.J., Xiao, B., Ha, Y., and Skehel, J.J. H1 and H7 influenza haemagglutinin structures extend a structural classification of haemagglutinin subtypes. Virology, 2004. 325(2): p. 287–96.

17. Rota, P.A., Wallis, T.R., Harmon, M.W., Rota, J.S., Kendal, A.P., and Nerome, K. Cocirculation of two distinct evolutionary lineages of influenza type B virus since 1983. Virology, 1990. 175(1): p. 59–68.

18. Benton, D.J., Nans, A., Calder, L.J., Turner, J., Neu, U., Lin, Y.P., Ketelaars, E., Kallewaard, N.L., Corti, D., Lanzavecchia, A., Gamblin, S.J., Rosenthal, P.B., and Skehel, J.J. Influenza hemagglutinin membrane anchor. Proceedings of the National Academy of Sciences, 2018. 115: p. 10112–10117.

19. Wilson, I.A., Skehel, J.J., and Wiley, D.C. Structure of the haemagglutinin membrane glycoprotein of influenza virus at 3 A resolution. Nature, 1981. 289(5796): p. 366–73.

20. Kirkpatrick, E., Qiu, X., Wilson, P.C., Bahl, J., and Krammer, F. The influenza virus hemagglutinin head evolves faster than the stalk domain. Sci Rep, 2018. 8(1): p. 10432.

21. Shih, A.C.-C., Hsiao, T.-C., Ho, M.-S., and Li, W.-H. Simultaneous amino acid substitutions at antigenic sites drive influenza A hemagglutinin evolution. Proceedings of the National Academy of Sciences, 2007. 104: p. 6283–6288.

22. Bizebard, T., Gigant, B.t., Rigolet, P., Rasmussen, B., Diat, O., BÃ¶seckei, P., Wharton, S.A., Skehel, J.J., and Knossow, M. Structure of influenza virus haemagglutinin complexed with a neutralizing antibody. Nature, 1995. 376(6535): p. 92–94.

23. Gao, J., Couzens, L., Burke, D.F., Wan, H., Wilson, P., Memoli, M.J., Xu, X., Harvey, R., Wrammert, J., Ahmed, R., Taubenberger, J.K., Smith, D.J., Fouchier, R.A.M., and Eichelberger, M.C. Antigenic Drift of the Influenza A(H1N1)pdm09 Virus Neuraminidase Results in Reduced Effectiveness of A/California/7/2009 (H1N1pdm09)-Specific Antibodies. MBio, 2019. 10(2).

24. Kasson, P.M. and Pande, V.S. Combining mutual information with structural analysis to screen for functionally important residues in influenza hemagglutinin. Pacific Symposium on Biocomputing. Pacific Symposium on Biocomputing, 2009: p. 492–503.

25. Popova, L., Smith, K., West, A.H., Wilson, P.C., James, J.A., Thompson, L.F., and Air, G.M. Immunodominance of Antigenic Site B over Site A of Hemagglutinin of Recent H3N2 Influenza Viruses. PLOS ONE, 2012. 7(7): p. e41895.

26. Wiley, D.C., Wilson, I.A., and Skehel, J.J. Structural identification of the antibody-binding sites of Hong Kong influenza haemagglutinin and their involvement in antigenic variation. Nature, 1981. 289(5796): p. 373–8.

27. Wilson, J.R., Guo, Z., Tzeng, W.-P., Garten, R.J., Xiyan, X., Blanchard, E.G., Blanchfield, K., Stevens, J., Katz, J.M., and York, I.A. Diverse antigenic site targeting of influenza hemagglutinin in the murine antibody recall response to A(H1N1)pdm09 virus. Virology, 2015. 485: p. 252–262.

28. Johnson, N.P. and Mueller, J. Updating the accounts: global mortality of the 1918-1920 “ Spanish” influenza pandemic. Bull Hist Med, 2002. 76(1): p. 105–15.

29. Lee, V.J., Chen, M.I., Chan, S.P., Wong, C.S., Cutter, J., Goh, K.T., and Tambyah, P.A. Influenza pandemics in Singapore, a tropical, globally connected city. Emerging infectious diseases, 2007. 13(7): p. 1052–1057.

30. Taubenberger, J.K. and Morens, D.M. Influenza: The Once and Future Pandemic. Public Health Reports, 2010. 125.

31. Viboud, C., Simonsen, L., Fuentes, R., Flores, J., Miller, M.A., and Chowell, G. Global Mortality Impact of the 1957-1959 Influenza Pandemic. J Infect Dis, 2016. 213(5): p. 738–45.

32. Dalby, A.R. and Iqbal, M. The European and Japanese outbreaks of H5N8 derive from a single source population providing evidence for the dispersal along the long distance bird migratory flyways. PeerJ, 2015. 3: p. e934.

33. Isoda, N., Twabela, A.T., Bazarragchaa, E., Ogasawara, K., Hayashi, H., Wang, Z.-J., Kobayashi, D., Watanabe, Y., Saito, K., Kida, H., and Sakoda, Y. Re-Invasion of H5N8 High Pathogenicity Avian Influenza Virus Clade 2.3.4.4b in Hokkaido, Japan, 2020. Viruses, 2020. 12(12).

34. Lewis, N.S., Banyard, A.C., Whittard, E., Karibayev, T., Al Kafagi, T., Chvala, I., Byrne, A., Meruyert, S., King, J., Harder, T., Grund, C., Essen, S., Reid, S.M., Brouwer, A., Zinyakov, N.G., Tegzhanov, A., Irza, V., Pohlmann, A., Beer, M., Fouchier, R.A.M., Akhmetzhan, S., and Brown, I.H. Emergence and spread of novel H5N8, H5N5 and H5N1 clade 2.3.4.4 highly pathogenic avian influenza in 2020. Emerging Microbes & Infections, 2021. 10(1): p. 148–151.

35. Vigeveno, R.M., Poen, M.J., Parker, E., Holwerda, M., de Haan, K., van Montfort, T., Lewis, N.S., Russell, C.A., Fouchier, R.A.M., de Jong, M.D., and Eggink, D. Outbreak Severity of Highly Pathogenic Avian Influenza A(H5N8) Viruses Is Inversely Correlated to Polymerase Complex Activity and Interferon Induction. Journal of Virology, 2020. 94.

36. Shi, J., Deng, G., Kong, H., Gu, C., Ma, S., Yin, X., Zeng, X., Cui, P., Chen, Y., Yang, H., Wan, X., Wang, X., Liu, L., Chen, P., Jiang, Y., Liu, J., Guan, Y., Suzuki, Y., Li, M., Qu, Z., Guan, L., Zang, J., Gu, W., Han, S., Song, Y., Hu, Y., Wang, Z., Gu, L., Yang, W., Liang, L., Bao, H., Tian, G., Li, Y., Qiao, C., Jiang, L., Li, C., Bu, Z., and Chen, H. H7N9 virulent mutants detected in chickens in China pose an increased threat to humans. Cell Res, 2017. 27(12): p. 1409–1421.

37. Subbarao, K. Avian influenza H7N9 viruses: a rare second warning. Cell Res, 2018. 28(1): p. 1–2.

38. Zhou, L., Chen, E., Bao, C., Xiang, N., Wu, J., Wu, S., Shi, J., Wang, X., Zheng, Y., Zhang, Y., Ren, R., Greene, C.M., Havers, F., Iuliano, A.D., Song, Y., Li, C., Chen, T., Wang, Y., Li, D., Ni, D., Zhang, Y., Feng, Z., Uyeki, T.M., and Li, Q. Clusters of Human Infection and Human-to-Human Transmission of Avian Influenza A(H7N9) Virus, 2013-2017. Emerg Infect Dis, 2018. 24(2).

39. Nachbagauer, R., Feser, J., Naficy, A., Bernstein, D.I., Guptill, J., Walter, E.B., Berlanda-Scorza, F., Stadlbauer, D., Wilson, P.C., Aydillo, T., Behzadi, M.A., Bhavsar, D., Bliss, C., Capuano, C., Carreño, J.M., Chromikova, V., Claeys, C., Coughlan, L., Freyn, A.W., Gast, C., Javier, A., Jiang, K., Mariottini, C., McMahon, M., McNeal, M., Solórzano, A., Strohmeier, S., Sun, W., Van der Wielen, M., Innis, B.L., García-Sastre, A., Palese, P., and Krammer, F. A chimeric hemagglutinin-based universal influenza virus vaccine approach induces broad and long-lasting immunity in a randomized, placebo-controlled phase I trial. Nature Medicine, 2021. 27(1): p. 106–114.

40. Beran, J., Peeters, M., Dewé, W., Raupachová, J., Hobzová, L., and Devaster, J.-M. Immunogenicity and safety of quadrivalent versus trivalent inactivated influenza vaccine: a randomized, controlled trial in adults. BMC Infectious Diseases, 2013. 13(1): p. 224.

41. Pépin, S., Donazzolo, Y., Jambrecina, A., Salamand, C., and Saville, M. Safety and immunogenicity of a quadrivalent inactivated influenza vaccine in adults. Vaccine, 2013. 31(47): p. 5572–5578.

42. Estrada, L.D. and Schultz-Cherry, S. Development of a Universal Influenza Vaccine. The Journal of Immunology, 2019. 202: p. 392–398.

43. Subbarao, K. and Matsuoka, Y. The prospects and challenges of universal vaccines for influenza. Trends in microbiology, 2013. 21(7): p. 350–358.

44. Chatziprodromidou, I.P., Arvanitidou, M., Guitian, J., Apostolou, T., Vantarakis, G., and Vantarakis, A. Global avian influenza outbreaks 2010-2016: a systematic review of their distribution, avian species and virus subtype. Syst Rev, 2018. 7(1): p. 17.

45. Del Rosario, J.M.M., Smith, M., Zaki, K., Risley, P., Temperton, N., Engelhardt, O.G., Collins, M., Takeuchi, Y., and Hufton, S.E. Protection From Influenza by Intramuscular Gene Vector Delivery of a Broadly Neutralizing Nanobody Does Not Depend on Antibody Dependent Cellular Cytotoxicity. Frontiers in Immunology, 2020. 11: p. 627.

46. Kanjilal, S. and Mina, M.J. Passive immunity for the treatment of influenza: quality not quantity. The Lancet Respiratory Medicine, 2019. 7(11): p. 922–923.

47. Manenti, A., Maciola, A.K., Trombetta, C.M., Kistner, O., Casa, E., Hyseni, I., Razzano, I., Torelli, A., and Montomoli, E. Influenza Anti-Stalk Antibodies: Development of a New Method for the Evaluation of the Immune Responses to Universal Vaccine. Vaccines, 2020. 8(1): p. 43.

48. Uyeki, T.M., Bernstein, H.H., Bradley, J.S., Englund, J.A., File, T.M., Fry, A.M., Gravenstein, S., Hayden, F.G., Harper, S.A., Hirshon, J.M., Ison, M.G., Johnston, B.L., Knight, S.L., McGeer, A., Riley, L.E., Wolfe, C.R., Alexander, P.E., and Pavia, A.T. Clinical Practice Guidelines by the Infectious Diseases Society of America: 2018 Update on Diagnosis, Treatment, Chemoprophylaxis, and Institutional Outbreak Management of Seasonal Influenzaa. Clin Infect Dis, 2019. 68(6): p. e1–e47.

49. WHO Global Influenza Surveillance Network. Manual for the Laboratory Diagnosis and Virological Surveillance of Influenza. 2011; Available from: http://www.who.int/influenza/gisrs_laboratory/manual_diagnosis_surveillance_influenza/en/index.html.

50. World Health Organization. Evaluation of influenza vaccine effectiveness: a guide to the design and interpretation of observational studies. 2017.

51. Hobson, D., Curry, R.L., Beare, A.S., and Ward-Gardner, A. The role of serum haemagglutination-inhibiting antibody in protection against challenge infection with influenza A2 and B viruses. The Journal of hygiene, 1972. 70(4): p. 767–777.

52. Alladi, C.S.H., Jagadesh, A., Prabhu, S.G., and Arunkumar, G. Hemagglutination Inhibition Antibody Response Following Influenza A(H1N1)pdm09 Virus Natural Infection: A Cross-Sectional Study from Thirthahalli, Karnataka, India. Viral Immunol, 2019. 32(5): p. 230–233.

53. Cox, R.J. Correlates of protection to influenza virus, where do we go from here? Hum Vaccin Immunother, 2013. 9(2): p. 405–8.

54. Agor, J.K. and Özaltın, O.Y. Models for predicting the evolution of influenza to inform vaccine strain selection. Human vaccines & immunotherapeutics, 2018. 14(3): p. 678–683.

55. Carnell, G.W., Trombetta, C.M., Ferrara, F., Montomoli, E., and Temperton, N.J. Correlation of Influenza B Haemagglutination Inhibiton, Single-Radial Haemolysis and Pseudotype-Based Microneutralisation Assays for Immunogenicity Testing of Seasonal Vaccines. Vaccines, 2021. 9(2).

56. Wallerström, S., Lagerqvist, N., Temperton, N.J., Cassmer, M., Moreno, A., Karlsson, M., Leijon, M., Lundkvist, Å., and Falk, K.I. Detection of antibodies against H5 and H7 strains in birds: evaluation of influenza pseudovirus particle neutralization tests. Infection Ecology & Epidemiology, 2014. 4(1): p. 23011.

57. Carnell, G.W., Ferrara, F., Grehan, K., Thompson, C.P., and Temperton, N.J. Pseudotype-Based Neutralization Assays for Influenza: A Systematic Analysis. Frontiers in Immunology, 2015. 6(161).

58. Temperton, N.J., Hoschler, K., Major, D., Nicolson, C., Manvell, R., Hien, V.M., Ha, D.Q., De Jong, M., Zambon, M., Takeuchi, Y., and Weiss, R.A. A sensitive retroviral pseudotype assay for influenza H5N1-neutralizing antibodies. Influenza and Other Respiratory Viruses, 2007. 1(3): p. 105–112.

59. Toon, K., Bentley, E.M., and Mattiuzzo, G. More Than Just Gene Therapy Vectors: Lentiviral Vector Pseudotypes for Serological Investigation. Viruses, 2021. 13(2).

60. Naldini, L., Blömer, U., Gallay, P., Ory, D., Mulligan, R., Gage, F.H., Verma, I.M., and Trono, D. In Vivo Gene Delivery and Stable Transduction of Nondividing Cells by a Lentiviral Vector. Science, 1996. 272: p. 263–267.

61. Zufferey, R., Nagy, D., Mandel, R.J., Naldini, L., and Trono, D. Multiply attenuated lentiviral vector achieves efficient gene delivery in vivo. Nature Biotechnology, 1997. 15(9): p. 871–875.

62. World Health Organization Global influenza strategy 2019-2030. 2019.

63. Cox, R.J., Mykkeltvedt, E., Robertson, J., and Haaheim, L.R. Non-Lethal Viral Challenge of Influenza Haemagglutinin and Nucleoprotein DNA Vaccinated Mice Results in Reduced Viral Replication. Scandinavian Journal of Immunology, 2002. 55(1): p. 14–23.

64. Raab, D., Graf, M., Notka, F., Schödl, T., and Wagner, R. The GeneOptimizer Algorithm: using a sliding window approach to cope with the vast sequence space in multiparameter DNA sequence optimization. Systems and Synthetic Biology, 2010. 4(3): p. 215–225.

65. Böttcher, E., Matrosovich, T., Beyerle, M., Klenk, H.-D., Garten, W., and Matrosovich, M. Proteolytic Activation of Influenza Viruses by Serine Proteases TMPRSS2 and HAT from Human Airway Epithelium. Journal of Virology, 2006. 80: p. 9896–9898.

66. Bertram, S., Glowacka, I., Blazejewska, P., Soilleux, E., Allen, P., Danisch, S., Steffen, I., Choi, S.-Y., Park, Y., Schneider, H., Schughart, K., and Pöhlmann, S. TMPRSS2 and TMPRSS4 facilitate trypsin-independent spread of influenza virus in Caco-2 cells. Journal of virology, 2010. 84(19): p. 10016–10025.

67. Ferrara, F. and Temperton, N. Pseudotype Neutralization Assays: From Laboratory Bench to Data Analysis. Methods and Protocols, 2018. 1(1): p. 8.

68. Miller, M., Pfeiffer, W., and Schwartz, T. Creating the CIPRES Science Gateway for inference of large phylogenetic trees. 2010 Gateway Computing Environments Workshop (GCE), 2010: p. 1–8.

69. Zhang, Y., Aevermann, B.D., Anderson, T.K., Burke, D.F., Dauphin, G., Gu, Z., He, S., Kumar, S., Larsen, C.N., Lee, A.J., Li, X., Macken, C., Mahaffey, C., Pickett, B.E., Reardon, B., Smith, T., Stewart, L., Suloway, C., Sun, G., Tong, L., Vincent, A.L., Walters, B., Zaremba, S., Zhao, H., Zhou, L., Zmasek, C., Klem, E.B., and Scheuermann, R.H. Influenza Research Database: An integrated bioinformatics resource for influenza virus research. Nucleic Acids Research, 2016. 45(D1): p. D466–D474.

70. Bosch, F.X., Garten, W., Klenk, H.D., and Rott, R. Proteolytic Cleavage of Influenza Virus Hemagglutinins: Primary Structure of the Connecting Peptide between HA, and HA2 Determines Proteolytic Cleavability and Pathogenicity of Avian Influenza Viruses. Virology, 1981. 113: p. 725–735.

71. Böttcher-Friebertshäuser, E., Garten, W., Matrosovich, M., and Klenk, H.D. The Hemagglutinin: A Determinant of Pathogenicity, in Influenza Pathogenesis and Control, Compans, R.W.and Oldstone, M.B.A., Editors. 2014, Springer.

72. Garten, W. and Klenk, H.D. Understanding Influenza Virus Pathogenicity. Trends in Microbiology, 1999. 7(3).

73. Ferrara, F., Molesti, E., Böttcher-Friebertshäuser, E., Cattoli, G., Corti, D., Scott, S.D., and Temperton, N.J. The human Transmembrane Protease Serine 2 is necessary for the production of Group 2 influenza A virus pseudotypes. Journal of molecular and genetic medicine : an international journal of biomedical research, 2012. 7: p. 309–314.

74. World Health Organization/World Organisation for Animal, H.F. and Agriculture Organization, H.N.E.W.G. Revised and updated nomenclature for highly pathogenic avian influenza A (H5N1) viruses. Influenza and Other Respiratory Viruses, 2014. 8(3): p. 384–388.

75. World Health Organization, Avian influenza: assessing the pandemic threat. 2005.

76. Petrova, V.N. and Russell, C.A. The evolution of seasonal influenza viruses. Nat Rev Microbiol, 2018. 16(1): p. 47–60.

77. Krammer, F. The human antibody response to influenza A virus infection and vaccination. Nature Reviews Immunology, 2019. 19(6): p. 383–397.

78. Krammer, F. and Palese, P. Advances in the development of influenza virus vaccines. Nat Rev Drug Discov, 2015. 14(3): p. 167–82.

79. Dreyfus, C., Laursen, N.S., Kwaks, T., Zuijdgeest, D., Khayat, R., Ekiert, D.C., Lee, J.H., Metlagel, Z., Bujny, M.V., Jongeneelen, M., van der Vlugt, R., Lamrani, M., Korse, H.J., Geelen, E., Sahin, O., Sieuwerts, M., Brakenhoff, J.P., Vogels, R., Li, O.T., Poon, L.L., Peiris, M., Koudstaal, W., Ward, A.B., Wilson, I.A., Goudsmit, J., and Friesen, R.H. Highly conserved protective epitopes on influenza B viruses. Science, 2012. 337(6100): p. 1343–8.

80. Corti, D., Voss, J., Gamblin, S.J., Codoni, G., Macagno, A., Jarrossay, D., Vachieri, S.G., Pinna, D., Minola, A., Vanzetta, F., Silacci, C., Fernandez-Rodriguez, B.M., Agatic, G., Bianchi, S., Giacchetto-Sasselli, I., Calder, L., Sallusto, F., Collins, P., Haire, L.F., Temperton, N., Langedijk, J.P., Skehel, J.J., and Lanzavecchia, A. A Neutralizing Antibody Selected from Plasma Cells That Binds to Group 1 and Group 2 Influenza A Hemagglutinins. Science, 2011. 333(6044): p. 850–6.

81. Mather, S.T., Wright, E., Scott, S.D., and Temperton, N.J. Lyophilisation of influenza, rabies and Marburg lentiviral pseudotype viruses for the development and distribution of a neutralisation-assay-based diagnostic kit. Journal of Virological Methods, 2014. 210: p. 51–58.

82. Ekiert, D., Bhabha, G., Elsliger, M.A., Friesen, R.H., Jongeneelen, M., Throsby, M., Goudsmit, J., and Wilson, I. Antibody Recognition of a Highly Conserved Influenza Virus Epitope. Science, 2009. 324(5924): p. 246–51.

83. Sui, J., Hwang, W.C., Perez, S., Wei, G., Aird, D., Chen, L.M., Santelli, E., Stec, B., Cadwell, G., Ali, M., Wan, H., Murakami, A., Yammanuru, A., Han, T., Cox, N.J., Bankston, L.A., Donis, R.O., Liddington, R.C., and Marasco, W.A. Structural and functional bases for broad-spectrum neutralization of avian and human influenza A viruses. Nat Struct Mol Biol, 2009. 16(3): p. 265–73.

84. Throsby, M., van den Brink, E., Jongeneelen, M., Poon, L.L., Alard, P., Cornelissen, L., Bakker, A., Cox, F., van Deventer, E., Guan, Y., Cinatl, J., ter Meulen, J., Lasters, I., Carsetti, R., Peiris, M., de Kruif, J., and Goudsmit, J. Heterosubtypic neutralizing monoclonal antibodies cross-protective against H5N1 and H1N1 recovered from human IgM+ memory B cells. PLoS One, 2008. 3(12): p. e3942.

85. Laursen, N.S. and Wilson, I.A. Broadly neutralizing antibodies against influenza viruses. Antiviral Res, 2013. 98(3): p. 476–83.

