## Supplementary Table 1 for "Establishment of pan-Influenza A (H1-H18) and pan-Influenza B (pre-split, Vic/Yam) Pseudotype Libraries for efficient vaccine antigen selection"

**Supplementary Table 1.** List of influenza hemagglutinin pseudotypes (PV) available at the Viral Pseudotype Unit, University of Kent. Protease employed to achieve the highest pseudotype titers are indicated.

| GROUP I INFLUENZA A HEMAGGLUTININ | | | | |
|---|---|---|---|---|
| **SUBTYPE** | **STRAIN** | **ACCESSION #** | **PLASMID** | **PROTEASE** |
| H1 | A/South Carolina/1/1918 | AF117241.1 | phCMV1 | TMPRSS4 |
| | A/Puerto Rico/8/1934 | AF389118.1 | pI.18 | TMPRSS4 |
| | A/New Caledonia/20/1999 | EU103824.1 | phCMV1 | TMPRSS4 |
| | A/duck/Italy/1447/2005 | HF563054.1 | pI.18 | TMPRSS4 |
| | A/Solomon Islands/3/2006 | EU124177.1 | pI.18 | TMPRSS4 |
| | A/Brisbane/59/2007 | CY163864.1 | pI.18 | TMPRSS4 |
| | A/California/7/2009 | CY121680.1 | pI.18 | TMPRSS4 |
| | A/Texas/05/2009 | GQ457487.1 | pI.18 | HAT |
| | A/England/195/2009 | GQ166661.1 | pEVAC | TMPRSS4 |
| | A/Bolivia/559/2013 | EPI466837 | pEVAC | TMPRSS4 |
| | A/swine/Guangxi/1/2013 2013_12_26_4 | KJ725056 | pEVAC | TMPRSS4 |
| | A/Michigan/45/2015 | EPI662594 | pEVAC | TMPRSS4 |
| | A/Slovenia/2903/2015 | EPI768541 | pEVAC | TMPRSS4 |
| | A/Brisbane/02/2018 | EPI1383389 | pEVAC | TMPRSS4 |
| | A/swine/Henan/SN10/2018 2018_02__4 | MN416619 | pEVAC | TMPRSS4 |
| | A/swine/Beijing/0301/2018 2018_03__4 | MN416589 | pEVAC | TMPRSS4 |
| H2 | A/Korea/426/1968 | CY125846.1 | pI.18 | HAT |
| | A/quail/Rhode Island/16-018622-1/2016 (H2) | KY272859 | pEVAC | TMPRSS4 |
| | A/duck/Germany/1215/1973 | CY014710.1 | pI.18 | TMPRSS4 |
| H5 | A/Hong Kong/156/1997 | AAC40508.1 | pI.18 | none |
| | A/Hong Kong/213/2003 | ABP51977.1 | pI.18 | none |
| | A/Vietnam/1194/2004 | ABP51976.1 | pI.18 | none |
| | A/Vietnam/1203/2004 | AB51977.1 | pI.18 | none |
| | A/Indonesia/5/2005 | ABW06108.1 | pI.18 | none |
| | A/turkey/Turkey/1/2005 | ABD73284.1 | pI.18 | none |
| | A/Anhui/1/2005 | ABD28180.1 | pI.18 | none |
| | A/whooper swan/Mongolia/244/2005 | GU186700.1<br>ACZ36881.1 | pEVAC<br>pI.18 | none |
| | A/bar-headed goose/Qinghai/2005 | BAE4815.1 | pI.18 | none |

| | | | | |
|---|---|---|---|---|
| | A/Jwe/Hong Kong/1038/2006 | ACJ26110.1A | pI.18 | none |
| | A/chicken/Mexico/07/2007 | KJ729343 | pEVAC | none |
| | A/Egypt/2629-NAMRU/2007 | ABM92273.1 | pI.18 | none |
| | A/chicken/Egypt 1709-01/2007 | ACD64996.1 | pI.18 | none |
| | A/chicken/Egypt 1709-06/2008 | ACD65000.1 | pI.18 | none |
| | A/gyrfalcon/Washington/41088-6/2014 | KP307984 | pI.18, pEVAC | none none |
| | A/mallard/Netherlands/41/2015 | MF694083 | pEVAC | none |
| H6 | A/American wigeon/California/HS007A/2015 | KY983173 | pEVAC | TMPRSS4 |
| | A/duck/Vietnam/HU9-455/2018 | LC497121 | pEVAC | TMPRSS4 |
| H8 | A/turkey/Ontario/6118/1968 | CY014659.1 | pI.18 | TMPRSS4 |
| | A/mallard duck/Netherlands/7/2015 | MF682649 | pEVAC | TMPRSS4 |
| | A/mallard duck/Ohio/16OS0672/2016 | MG280005 | pEVAC | TMPRSS4 |
| H9 | A/Hong Kong/1073/1999 | AJ404626.1 | pI.18 | TMPRSS4 |
| | A/chicken/Israel/291417/2017 | MH558944 | pEVAC | TMPRSS4 |
| H11 | A/duck/Memphis/546/1974 | AB292779 | phCMV1 | TMPRSS4 |
| | A/common teal/Netherlands/1/2015 | MF693986 | pEVAC | TMPRSS4 |
| | A/red shoveler/Chile/C14653/2016 | MH134837 | pEVAC | TMPRSS4 |
| H12 | A/duck/Alberta/60/1976 | CY130078.1 | phCMV1 | HAT |
| | A/duck/Mongolia/850/2018 | MK979051 | pEVAC | TMPRSS4 |
| | A/Northern Shoveler/Nevada/D1516557/2015 | MK928236 | pEVAC | TMPRSS4 |
| H13 | A/laughing gull/New Jersey/UGAI17-2843/2017 | MH068343 | pEVAC | TMPRSS4 |
| | A/ring-billed gull/Minnesota/OPMNAI0816/2017 | MH763859 | pEVAC | TMPRSS4 |
| H16 | A/black-headed gull/Sweden/2/1999 | AY684888.1 | phCMV1 | HAT |
| | A/black-headed gull/Netherlands/1/2016 | MF694134 | pEVAC | TMPRSS4 |
| | A/Mew Gull/Southcentral Alaska/18MB01898/2018 | MN210308 | pEVAC | TMPRSS4 |
| H17 | A/Little shouldered bat/Guatemala/060/2011 | CY103892.1 | pI.18 | HAT |
| H18 | A/flat-faced bat/Peru/33/2010 | CY125945 | pEVAC | T4 |

| GROUP II INFLUENZA A HEMAGGLUTININ | | | | |
|---|---|---|---|---|
| **SUBTYPE** | **STRAIN** | **ACCESSION #** | **PLASMID** | **PROTEASE** |
| H3 | A/Texas/50/2012 | KC892952.1 | pI.18 | TMPRSS4 |
| | A/Udorn/307/1972 | DQ508929.1 | pI.18 | TMPRSS2 |
| | A/California/7/2004 | CY114373.1 | pEVAC | HAT |
| | A/Wisconsin/67/2005 | CY034116.1 | pEVAC | HAT |
| | A/Japan/WRAIR1059P/2008 | | pEVAC | HAT |
| | A/Switzerland/9715293/2013 | EPI814528 | pEVAC | HAT |
| | A/New Caledonia/71/2014 | EPI551570 | pEVAC | HAT |
| | A/duck/Quang Ninh/220/2014 | LC053492 | pEVAC | HAT |
| | A/ruddy turnstone/Delaware Bay/606/2017 | MH135712 | pEVAC | TMPRSS2 |
| | A/Kansas/14/2017 | Vaccine strain, 3c3.A | pEVAC | HAT |
| | A/Switzerland/8060/2017 | EPI1326015 | pEVAC | TMPRSS4 |
| | A/South Australia/34/2019 | EPI1607117 | pEVAC | HAT |
| H4 | A/duck/Czechoslovakia/1956 | D90302.1 | phCMV1 | TMPRSS2 |
| | A/green-winged teal/California/K218/2005 | CY045351 | pEVAC | TMPRSS2 |
| | A/Calidris ruficollis/Hokkaido/12EY0172/2012 | LC467224 | pEVAC | TMPRSS2 |
| H7 | A/FPV/Rostock/1934 | AAA43150 | phCMV1 | none |
| | A/chicken/Pakistan/34668/1995 | CY035831 | pI.18 | none |
| | A/chicken/Italy/1082/1999 | CY022677 | pI.18 | TMPRSS2 |
| | A/chicken/Italy/13474/1999 | AJ491720 | pI.18 | none |
| | A/chicken/Netherlands/1/2003 | AAR02640.1 | pI.18 | none |
| | A/chicken/Netherlands/219/2003 | AAR02640.1 | pI.18 | none |
| | A/Shanghai/2/2013 | KF021597 EPI448936 | pI.18, pEVAC | TMPRSS4 |
| | A/northern pintail duck/California/UCD1582/2016 | MH251202 | pEVAC | TMPRSS4 |
| | A/duck/Viet Nam/HU10-64/2018 | MK629228 | pEVAC | TMPRSS4 |
| | A/Anhui/1/2013 | CY187618.1 | pEVAC | TMPRSS4 |
| H10 | A/duck/Bangladesh/24268/2015 | MH071504 | pEVAC | TMPRSS4 |
| | A/mallard/Utah/D1802334/2018 | MK995817 | pEVAC | TMPRSS4 |
| | A/chicken/Germany/N49 | CY014671.1 | pEVAC | TMPRSS4 |
| H14 | A/mallard/Astrakhan/263/1982 | AB289335.1 CY014604 | phCMV1 pEVAC | TMPRSS4 |

| | | | | |
|---|---|---|---|---|
| | A/blue-winged Teal/Ohio/18OS1695/2018 | MN431050 | pEVAC | TMPRSS4 |
| H15 | A/shearwater/West Australia/2576/1979 | CY130102.1<br>CY006010 | phCMV1<br>pEVAC | TMPRSS4 |
| | A/duck/Bangladesh/24697/2015 | KY635719 | pEVAC | TMPRSS4 |
| **INFLUENZA B** | | | | |
| **SUBTYPE** | **STRAIN** | **ACCESSION #** | **PLASMID** | **PROTEASE** |
| B | B/Hong Kong/8/1973 | K00425 | phCMV1 | HAT |
| | B/Victoria/2/1987 | FJ766840 | phCMV1 | HAT |
| | B/Yamagata/16/1988 | CY018765.1 | phCMV1 | HAT |
| | B/Florida/4/2006 | EU515992 | phCMV1 | HAT |
| | B/Bangladesh/3333/2007 | CY115255.1 | pI.18 | HAT |
| | B/Brisbane/60/2008 | KX058884.1<br>EPI753679 | pI.18<br>pEVAC | HAT |
| | B/Phuket/3073/2013 | | pEVAC | HAT |
| | B/Colorado/06/2017 | | pEVAC | HAT |
| | B/Washington/2/2019 | EPI1368874 | pEVAC | HAT |
